# Is zebrafish heart regeneration “complete”? Lineage-restricted cardiomyocytes proliferate to pre-injury numbers but some fail to differentiate in fibrotic hearts

**DOI:** 10.1101/2020.09.08.277939

**Authors:** Alberto Bertozzi, Chi-Chung Wu, Phong D. Nguyen, Mohankrishna Dalvoy Vasudevarao, Medhanie A. Mulaw, Charlotte D. Koopman, Teun P. de Boer, Jeroen Bakkers, Gilbert Weidinger

## Abstract

Adult zebrafish are frequently described to be able to “completely” regenerate the heart. Yet, the extent to which cardiomyocytes lost to injury are replaced is unknown, since only indirect or non-quantitative evidence for cardiomyocyte proliferation exists. We established stereological methods to quantify the number of cardiomyocytes at several time-points post cryoinjury. Intriguingly, after cryoinjuries that killed about 1/3 of the ventricular cardiomyocytes, pre-injury cardiomyocyte numbers were restored already within 30 days. Yet, many hearts retained small residual scars, and a subset of cardiomyocytes bordering these fibrotic areas remained smaller, lacked differentiated sarcomeric structures, and displayed defective calcium signaling. Thus, a subset of regenerated cardiomyocytes failed to fully mature. While lineage-tracing experiments have shown that regenerating cardiomyocytes are derived from differentiated cardiomyocytes, technical limitations have previously made it impossible to test whether cardiomyocyte trans-differentiation contributes to regeneration of non-myocyte cell lineages. Using Cre responder lines that are expressed in all major cell types of the heart, we found no evidence for cardiomyocyte transdifferentiation into endothelial, epicardial, fibroblast or immune cell lineages. Overall, our results imply a refined answer to the question whether zebrafish can completely regenerate the heart: in response to cryoinjury, preinjury cardiomyocyte numbers are indeed completely regenerated, while restoration of cardiomyocyte differentiation and function, as well as resorption of scar tissue, is less robustly achieved.

## Introduction

In adult mammals, heart injuries result in permanent scarring and lost cardiomyocytes cannot be regenerated. In contrast, zebrafish hearts can achieve scar-free healing and show signs of muscle restoration following several types of injuries (Chablais et al., 2011; Gonzalez-Rosa et al., 2011; Poss et al., 2002; Schnabel et al., 2011; Wang et al., 2011). In fact, zebrafish have frequently been described to be able to “completely” regenerate their hearts (Chablais et al., 2011; Gonzalez-Rosa et al., 2011). Arguably, one of the most important aspects of heart regeneration is the restoration of cardiomyocytes lost to injury. Cardiomyocytes show robust cell cycle activity in injured zebrafish hearts, and many studies have quantified this using PCNA immunofluorescence or nucleotide analog incorporation during S-phase (Chablais et al., 2011; Gonzalez-Rosa et al., 2011; Poss et al., 2002; Sallin et al., 2015; Schnabel et al., 2011; Wang et al., 2011; Wu et al., 2016). Unfortunately, more direct quantitative evidence for cardiomyocyte proliferation is largely lacking, mainly since phospho-Histone 3 as readout for cardiomyocyte mitosis is very rarely detected (0-2 pH3-positive cells per heart section even at the peak of cell cycle activity, A.B. and G.W., unpublished and (Sallin et al., 2015)) and thus is rarely used in zebrafish heart regeneration studies. Other existing evidence for cardiomyocyte regeneration is likewise indirect or qualitative, like the restoration of the area covered by muscle (Poss et al., 2002), and the expansion of genetically labelled clones of cardiomyocytes (Kikuchi et al., 2010). In summary, the number of regenerated cardiomyocytes has not been quantified and it is unknown whether zebrafish hearts regenerate all cardiomyocytes lost to injury. Of note, it has been shown that a thin layer of cardiomyocytes, which are located subcortically and have been termed “primordial”, does not completely regenerate (Pfefferli and Jazwinska, 2017).

Injured hearts form fibrin- and collagen-rich wound tissue, in particular after cryoinjury, which is largely resolved during later stages of regeneration (Chablais and Jazwinska, 2012). Thus, many studies have used quantification of wound size to assess regenerative success, often using the implicit assumption that resorbed wound tissue is being replaced by regenerated muscle. However, it is conceivable that wound resorption can occur in the absence of cardiomyocyte proliferation. One line of evidence that supports this is the observation that cardiomyocyte cell cycle activity peaks at 7 days post injury (dpi) (Chablais et al., 2011; Gonzalez-Rosa et al., 2011; Schnabel et al., 2011) and returns to basal levels at around 30 dpi (Chablais et al., 2011), while scar resorption has been reported to take 60 to 130 days in cryoinjured hearts (Chablais et al., 2011; Gonzalez-Rosa et al., 2011). This raises the question whether wound resorption is fully correlated with cardiomyocyte restoration, and whether it thus can be used as readout for cardiomyocyte regeneration. Finally, several aspects of heart function, which depend on cardiomyocyte activity, do not seem to fully recover in cryoinjured hearts, in particular ventricular wall motion (Gonzalez-Rosa et al., 2014; Gonzalez-Rosa et al., 2011; Hein et al., 2015; Wang et al., 2011).

Overall, we think there is a need to clarify whether zebrafish can indeed “completely” regenerate the heart. In particular, the extent to which zebrafish cardiomyocytes regenerate should be quantified, given the wide-spread use of the model. Here, we have used cardiomyocyte counting to quantify the true extent of zebrafish cardiomyocyte restoration.

Fate-mapping studies have shown that new cardiomyocytes arise from differentiated spared cardiomyocytes during heart regeneration (Gupta et al., 2013; Gupta and Poss, 2012; Jopling et al., 2010; Kikuchi et al., 2010). Differentiated cells are increasingly being recognized as in important source of repair cells also in other systems (Merrell and Stanger, 2016; Mills et al., 2019; Tanaka, 2016; Tata and Rajagopal, 2016; Zhang and Liu, 2020). In many cases, differentiated cells dedifferentiate, that is they lose aspects of their mature state, but remain lineage-restricted. Yet, several intriguing cases of transdifferentiation, where cells give rise to other cell types in the course of regeneration, have also been reported (Merrell and Stanger, 2016; Tata and Rajagopal, 2016; Zhang and Liu, 2020). Thus, the demands of regeneration can alter the fate and potency of adult cells. Whether cardiomyocytes acquire multipotency and give rise to other cell types during zebrafish heart regeneration has not been addressed, since all lineage tracing studies used transgenic lines expressing cardiomyocyte-specific Cre recombinase in combination with Cre-responding tracer lines which are only expressed in cardiomyocytes (Gupta et al., 2013; Gupta and Poss, 2012; Jopling et al., 2010; Kikuchi et al., 2010). These existing tools are therefore not suitable for detecting contribution of cardiomyocytes to other cell types.

Here, using a quantitative approach, we find that zebrafish cardiomyocyte numbers are restored to pre-injury levels within 30 days of injury. However, persistent fibrosis appears to interfere with cardiomyocyte differentiation, resulting in incomplete maturation of a subset of regenerated cardiomyocytes in hearts that retain scars. Using newly created lineage-tracing tools, we find no evidence of cardiomyocyte contribution to other major cardiac cell types. Our results reveal the enormous potency of zebrafish hearts to completely regenerate all cardiomyocytes lost to injury via proliferation of lineage-restricted cells.

## Methods

Expanded Materials & Methods can be found in the Supplemental Materials. The raw data that support the findings of this study are available from the corresponding author upon reasonable request.

### Transgenic fish lines and cryoinjury

All procedures involving animals were approved by the state of Baden-Württemberg (Project numbers 1352 and 1427) and by local animal experiment committees.

Zebrafish (*Danio rerio*) of ∼ 4 to 6 months of age were used, except for experiments with juvenile fish, which ranged from standardized standard length 11 to 16 mm (∼ 8 weeks of age in our facility). The following previously published transgenic fish lines were used: *fli1a*:eGFP^y1Tg^ (Lawson and Weinstein, 2002), Et(*krt4*:eGFP)^sqet27^ (Parinov et al., 2004), *coro1a*:eGFP^hkz04tTg^ (Li et al., 2012), *postnb*:Citrine^cn6Tg^ (Sanchez-Iranzo et al., 2018), *wt1b*:EGFP^li1Tg^ (Perner et al., 2007), *cryaa*:DsRed,-5.1*myl7*:Cre-ERT2^pd10^ (Kikuchi et al., 2010), *hsp70l*:LOXP-DsRed2-LOXP-NLS-EGFP^tud9^ (Knopf et al., 2011), *myl7*:EGFP^twu34^ (Huang et al., 2003) and -5.1*myl7*:DsRed2-nls^f2T^ (Mably et al., 2003). Cryoinjuries and sham injuries were performed as previously described, with the exception that a liquid-nitrogen cooled copper filament was used to freeze the heart (Schnabel et al., 2011).

### Creation of hsp70l:LOXP-luc-myc-STOP-LOXP-dTomato, cryaa:YPet^ulm9^ (hsp:L to T) and myl7:GCaMP6f-nls-T2A-RCaMP107-NES-pA)^hu11799^ transgenic animals

Both lines were created using Tol2 mediated transgenesis. For the *hsp70l*:LOXP-luc-myc-STOP-LOXP-dTomato, *cryaa*:YPet^*ulm9*^ (*hsp*:L to T) line, the following elements were assembled by PCR: attP site, zebrafish *hsp70l* promoter (Halloran et al., 2000), loxP site, firefly luciferase fused at its C-terminus to a 5x myc-tag, ocean pout polyA STOP cassette (Clark et al., 2011), loxP site, dTomato, SV40 polyA, zebrafish *cryaa* promoter (Kurita et al., 2003), YPet, ocean pout polyA STOP cassette, attP site, AmCyan, SV40 polyA. The Tg(*myl7*:GCaMP6f-nls-T2A-RCaMP107-NES-pA)^hu11799^ was designed to report nuclear calcium transients in green using the fluorescent protein (GCaMP6f nls, Chen et al., 2013) and cytoplasmic calcium transients in red, using (RCaMP107 NES, Ohkura et al., 2012). However, in the transgenic line, the GCaMP6f nls protein was found to be located in the cytoplasm, not the nucleus. Thus, only cytosolic calcium transients were measured (red).

### Sectioning and imaging

Hearts were fixed in 4% PFA (in 0.1 M phosphate buffer pH 7.4 with 4% sucrose) at room temperature for 1 h, embedded in OCT medium and cryo-sectioned into 10 µm thick sections unless stated otherwise. For cardiomyocyte quantification experiments, hearts were embedded with a random orientation to ensure that no stereological bias was introduced in our analysis. Heart sections were equally distributed onto 6 serial slides or multiples thereof to accommodate sections representing all areas of the ventricle onto each slide. Images were acquired using an Olympus BX60 compound microscope (wound size), a Keyence microscope (widefield fluorescence for nuclei counting), or a Leica SP5 confocal microscope (all other data).

### Immunofluorescence and histological staining

EGFP and DsRed in *myl7*:EGFP^twu34^ and -5.1*myl7*:DsRed2-nls^f2T^ transgenic fish, respectively, was detected by imaging of native EGFP and DsRed fluorescence. All other markers were detected via immunofluorescence.

Primary antibodies used were anti-aldh1a2 (Abmart #P30011, 1:500), anti-RFP (Genetex #GTX82561, 1:300), anti-dsRed (Clontech #632496, 1:300), anti-c-Myc (Santa Cruz #SC40, 1:50), anti-PCNA (Dako #M0879, 1:1000), anti-GFP (Abcam #ab13970, 1:500), anti-MHC (Developmental Studies Hybridoma Bank, MF20, 1:50), anti-Myl7 (Genetex, #GTX128346, 1:300), anti-laminin (Sigma-Aldrich, #L9393, 1:200), anti-alpha-actinin (Sigma-Aldrich #A7811, 1:500), anti-MLCK (Sigma-Aldrich #M7905, 1:100), anti-Vimentin (Developmental Studies Hybridoma Bank #40E-C, 1:50), anti-Col1a1a (Developmental Studies Hybridoma Bank #SP1.D8, 1:50), anti-embMHC (Developmental Studies Hybridoma bank, # N2.261, 1:50) and anti-Mef2 (Santa Cruz #SC313, 1:50). Secondary antibodies conjugated to Alexa 488, 555 or 633 (Invitrogen) were used at a dilution of 1:1000. Nuclei were shown by DAPI (4’,6’-diamidino-2-phenylindole) staining. Immunostainings were performed as previously described (Wu et al., 2016). Acid fuchsin orange G (AFOG) staining was performed as previously described (Poss et al., 2002).

### Cardiomyocyte lineage tracing

For adult experiments, *cryaa*:DsRed,-5.1*myl7*:Cre-ERT2^pd10^; *hsp70l*:LOXP-luc-myc-STOP-LOXP-dTomato, *cryaa*:YPet^ulm9^ double transgenic fish were injected intraperitoneally with 20 µl of 1.3 mM 4-OH Tamoxifen (4-OHT) in dH_2_O once daily for 6 days. Fish were heat-shocked once daily at 37^°^ C for 1 h, after which the water temperature was reduced back to 27^°^ C within 15 minutes, for 6 days before harvesting.

For juvenile experiments, *cryaa:*DsRed,-5.1*myl7*:Cre-ERT2^pd10^; *hsp70l*:LOXP-DsRed2-LOXP-NLS-EGFP^tud9^ double transgenic embryos were incubated with 10 µM 4-OHT diluted in E3 medium at 28.5^°^ C from 1 to 4 days post fertilization (dpf). Fresh 4-OHT solution was applied daily. Embryos with successful cardiomyocyte labelling were selected at 5 dpf after a single heat-shock by GFP fluorescence and raised until juvenile stages (8 weeks). Fish were heat-shocked once daily at 37^°^ C for 1 h for 2 days before harvesting.

### Statistical analyses

p-values are reported in figures, and information about samples sizes and types of statistical tests used can be found in figure legends. p-values larger than 0.01 are reported to two decimal places, those between 0.01 and 0.001 to three decimal places, and p-values smaller than 0.001 are reported as <0.001. Error bars represent CI 95%, except in Fig. 5 and 6, where they represent s.e.m. To support statements about the lack of differences between experimental groups (where p-values are > 0.05) we computed the effect size (based on the Means, SDs and sample sizes of the experimental groups), and from this the smallest significant differences that we could have detected. These and the observed differences are reported in the Figure legends. If we considered the smallest significant detectable difference as biologically not meaningful, we concluded that no difference exists.

## Results

### Cardiomyocyte cell cycle and wound resorption dynamics do not fully correlate

Wound size and cardiomyocyte cell cycle activity are commonly used as readouts for the regenerative response of zebrafish hearts. Using PCNA immunostaining and EdU incorporation, we found that cardiomyocyte cell cycle activity in cryoinjured hearts peaked at 7 dpi, dropped dramatically by 14 dpi and was only marginally higher than in uninjured hearts by 30 dpi (Figure 1A, B, Supplementary Figure 1A, B), consistent with a previous report (Bednarek et al., 2015). However, wound tissue was still observed in 65% of hearts at 30 dpi (Figure 1C, D). This apparent discrepancy between the dynamics of cardiomyocyte cell cycle activity and wound resorption could either indicate that wound size is not a good proxy for the extent of cardiomyocyte regeneration or that cardiomyocytes do not regenerate to pre-injury numbers.

**Figure 1.**
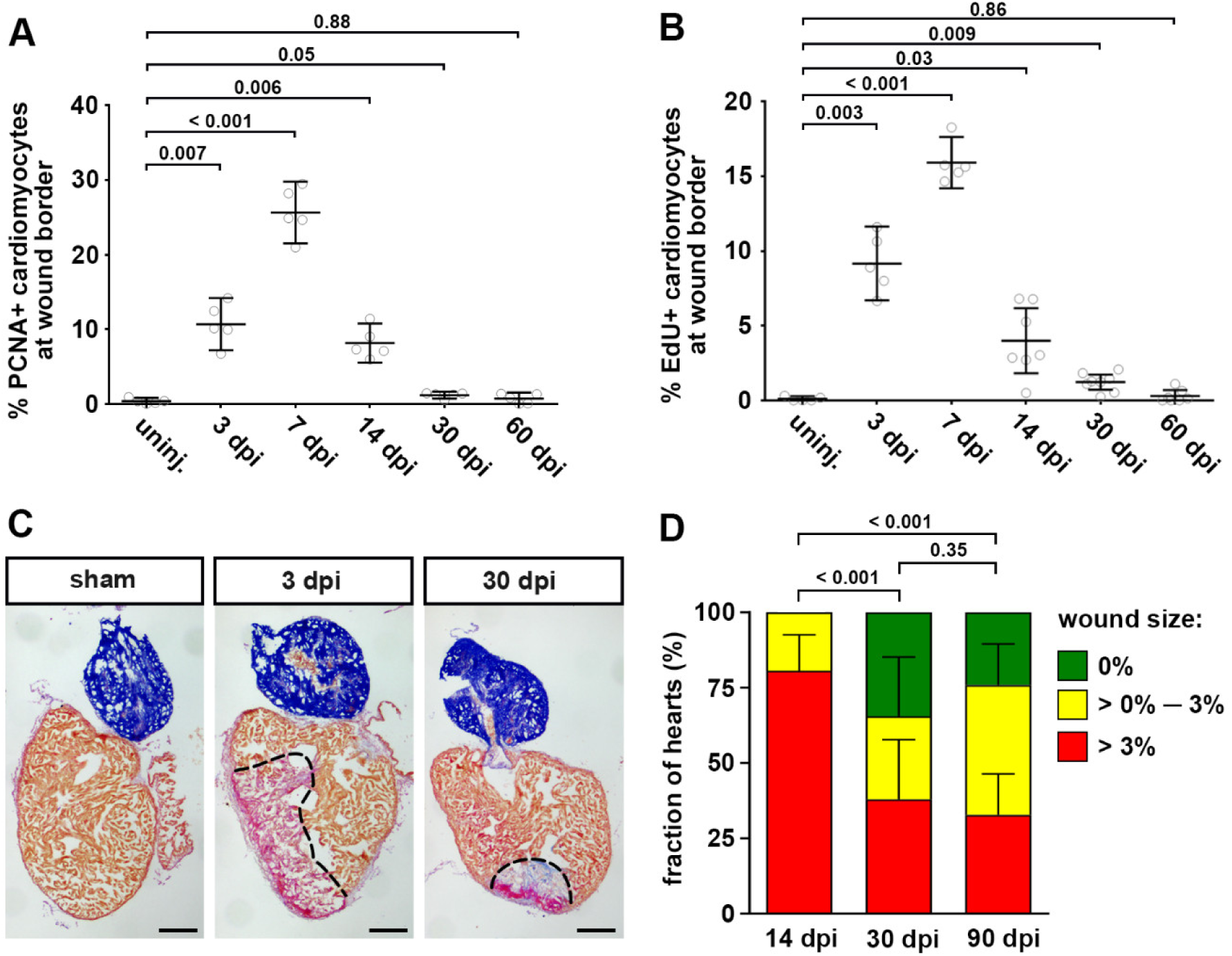
Cardiomyocyte cell cycle activity returns to basal levels prior to complete morphological regeneration. (A) The fraction of PCNA^+^ EGFP^+^ cardiomyocytes in *myl7*:EGFP^twu34^ fish within 150 µm of the wound border peaks at 7 dpi and drops to baseline by 30 dpi. uninj, uninjured. Error bars represent CI 95%. N = 5 hearts for each group. One-way ANOVA + Dunnett’s multiple comparisons test. Numbers indicate p-value. (B) Fraction of EdU^+^ EGFP^+^ cardiomyocytes in *myl7*:EGFP^twu34^ fish. For each time point, a single intraperitoneal injection of EdU was performed 24 hours prior to heart extraction. n (hearts) = 5 (uninjured), 5 (3 dpi), 5 (7 dpi), 7 (14 dpi), 8 (30 dpi), 7 (60 dpi). One-way ANOVA + Dunnett’s multiple comparisons test. (C) Many hearts contain wound tissue detected by Acid Fuchsin Orange G (AFOG) staining at 30 dpi. Red: fibrin, blue: collagen, brown: myocardium. Scale bars, 100 µm. (D) Percentage of hearts without residual wounds (= 0% of ventricle area), small wounds (between 0% and 3%) or bigger wounds (> 3%) at 14, 30 and 90 dpi. n = 31 (14 dpi), 29 (30 dpi), 58 (90 dpi). Chi-square test.

### Cardiomyocyte numbers are fully restored during heart regeneration within 30 days

To quantify the extent of cardiomyocyte regeneration, we measured cardiomyocyte numbers by counting nuclei labeled by DsRed in -5.1*myl7*:DsRed2-nls^f2Tg^ transgenic fish, since ∼95% of adult zebrafish cardiomyocytes are mononucleated (Wills et al., 2008). We chose to avoid variability potentially introduced by heart dissociation and counted cardiomyocytes on heart sections, using stereological methods to reconstruct cardiomyocyte numbers in the three-dimensional heart volume from two-dimensional sections (Figure 2A). We found that semi-automated analysis of images of an evenly spaced subset of cryosections acquired by wide-field fluorescent microscopy represented a reliable method to determine cardiomyocyte numbers (Figure 2A, Supplementary Figure 2A-D, see Extended Methods in the Supplemental Material for details).

**Figure 2.**
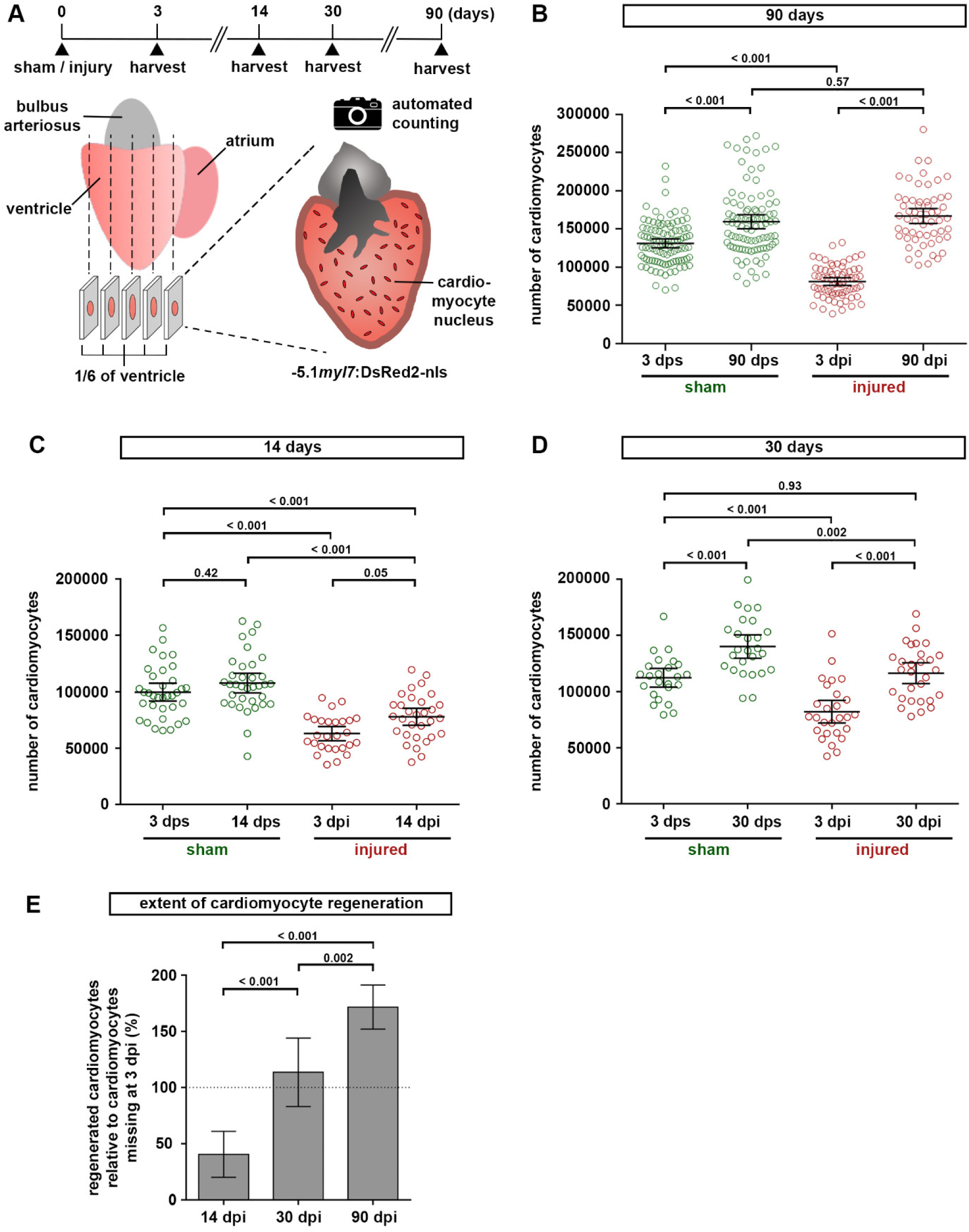
Zebrafish hearts regenerate all cardiomyocytes lost to injury. (A) Experimental design for quantification of cardiomyocytes. (B) Cardiomyocytes regenerate to the same number as found in sham injured hearts within 90 days. The total number of ventricular cardiomyocytes per heart is plotted. Error bars, CI 95%. n (hearts) = 99 (3 dps), 92 (90 dps), 68 (3 dpi), 58 (90 dpi). One-way ANOVA + Tukey’s multiple comparisons test. Observed difference 90 dpi vs 90 dps = 7400 (4% of CMs in 90 dps); smallest significant detectable difference = 19800 (12%). (C) Cardiomyocyte regeneration is not complete at 14 dpi. n (hearts) = 35 (3 dps), 35 (14 dps), 27 (3 dpi), 31 (14 dpi). One-way ANOVA + Tukey’s multiple comparisons test. (D) Pre-injury cardiomyocyte numbers have been restored at 30 dpi. n (hearts) = 24 (3 dps), 26 (30 dps), 27 (3 dpi), 29 (30 dpi). One-way ANOVA + Tukey’s multiple comparisons test. Observed difference 30 dpi vs 3 dps = 4100 (3.6% of CMs in 3 dps); smallest significant detectable difference = 17600 (15%). (E) The number of regenerated cardiomyocytes at 14, 30 and 90 dpi relative to the number missing in the respective 3 dpi group is plotted. Dotted line represents complete regeneration. n (hearts) = 31 (14 dpi), 29 (30 dpi), 58 (90 dpi). One-way ANOVA + Tukey’s multiple comparisons test.

We randomly assigned -5.1*myl7*:DsRed2-nls^f2Tg^ transgenic individuals from a large cohort of fish to either sham injury or ventricle cryoinjury. Within each group, fish were randomly selected for analysis at 3 or 90 days post intervention (Figure 2A). Sham injured control ventricles contained on average 131,000 ± 5,000 cardiomyocytes at 3 days post sham injury (dps, mean ± 95% CI, Figure 2B). At 3 days post cryoinjury (dpi) wound area represented 30 ± 2% of the ventricle area, and 38 ± 5% of cardiomyocytes had been lost to injury (Figure 2B, Supplementary Figure 3A). In sham injured hearts, cardiomyocyte numbers and the ventricular volume had increased significantly by 90 dps, indicating that hearts grew by cardiomyocyte hyperplasia under our housing conditions (Figure 2B, Supplementary Figure 3B). Importantly, no significant difference in cardiomyocyte numbers was observed between sham and cryoinjured hearts at 90 days (Figure 2B), and ventricular volume had increased even beyond sham levels in injured hearts (Supplementary Figure 3B). We conclude that zebrafish hearts fully regenerate the pre-injury cardiomyocyte number. Intriguingly, at 90 dpi the number of cardiomyocytes was 172 ± 20% of the number of cardiomyocytes lost to injury (the number of cells missing at 3 dpi, Figure 2E). Thus, lost cardiomyocytes had not only been fully restored, but regenerated hearts had grown by hyperplasia to the same extent as sham injured hearts (Figure 2B).

To assess the time span required for complete cardiomyocyte regeneration, we counted cardiomyocytes at 14 and 30 days post sham or cryoinjury. At 14 dpi, 40 ± 20% of the cardiomyocytes lost to injury had been regenerated (Figure 2C, E). At 30 days post cryoinjury, cardiomyocyte numbers did not significantly differ from those in 3 dps hearts, showing that pre-injury cardiomyocyte numbers had already been restored (Figure 2D). Indeed, 114 ± 30% of the cardiomyocytes lost to injury had been regenerated (Figure 2E). However, cardiomyocyte numbers at 30 dpi were lower than those seen in sham injured hearts at 30 dps (Figure 2D), indicating that 30 days were just sufficient to complete cardiomyocyte regeneration, but not to accompany additional hyperplasia of the myocardium. These data show that regeneration of cardiomyocyte numbers is completed much earlier than what is commonly considered the end point for zebrafish heart regeneration after cryoinjury based on scar resolution (Chablais et al., 2011; Gonzalez-Rosa et al., 2011; Schnabel et al., 2011).

### Cardiomyocytes withdraw from the cell cycle in hearts displaying incomplete scar resorption

Although pre-injury cardiomyocyte numbers had been fully restored at 90 dpi, the majority of hearts (75%) still contained collagen-rich residual wounds, which accounted for > 3% of ventricular area in 33% of the hearts (Figure 3A, 1D). Interestingly, neither the average wound area nor the fraction of hearts retaining a wound > 3% of ventricle area changed between 30 dpi and 90 dpi (Figure 3B, 1D, Supplementary Figure 3A). This suggests that those hearts with residual wounds at 30 dpi might never achieve scar-free regeneration. Thus, we wondered whether in such hearts cell-cycling and regeneration of cardiomyocytes was still ongoing. However, the number of PCNA^+^ cardiomyocytes was equally at basal level in hearts with or without wounds (Figure 3C, D). Together, these data show that border-zone cardiomyocytes exit the cell cycle even when residual fibrotic tissue has not been completely resorbed.

**Figure 3.**
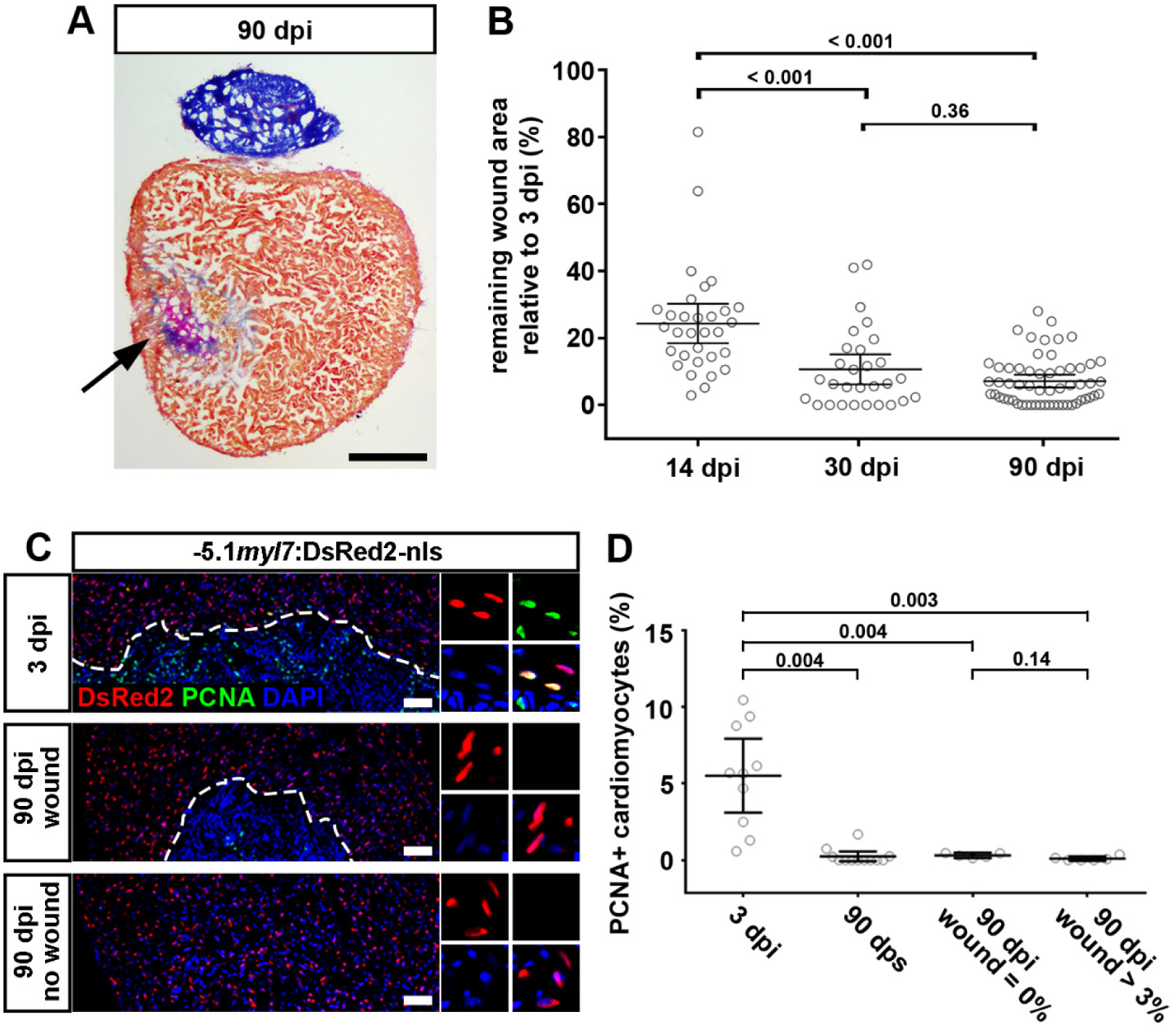
Cardiomyocyte proliferation ceases in both scarred and unscarred hearts. (A) AFOG stained section at 90 dpi displaying incomplete scar resorption (arrow). Scale bar, 100 µm. (B) Average wound area does not significantly shrink between 30 and 90 dpi. n (hearts) = 31 (14 dpi), 29 (30 dpi), 58 (90 dpi). One-way ANOVA + Tukey’s multiple comparisons test. Observed difference 90 dpi vs 30 dpi = 3.5 (32% of 30 dpi wound area); smallest significant detectable difference = 5.8 (54% of 30 dpi wound area). (C, D) PCNA+ cardiomyocytes can be detected at 3 dpi, but not in 90 dpi hearts with or without wound. Scale bar, 100 µm. n (hearts) = 10 (3 dpi), 12 (90 dps), 5 (90 dpi no wound), 6 (90 dpi wound). One-way ANOVA + Tukey’s multiple comparisons test.

### Regenerated cardiomyocytes remain small and fail to fully mature in areas enclosing residual scars

Since the majority of hearts retained some scar tissue in our experiments, we asked whether the overall morphology of the regenerated myocardium was altered. However, we observed no difference between sham and injured hearts in volume of the healthy myocardium, overall cardiomyocyte density, or thickness of the cortical layer identified by expression of laminin (Sanchez-Iranzo et al., 2018) (Supplementary Figure 4A-D). However, at 30 and 90 dpi, hearts that retained scars displayed a higher cardiomyocyte density in the external wound border zone than at the non-injured base (Figure 4A, B, Supplementary Figure 5A), consistent with a previous report (Gonzalez-Rosa et al., 2014). We wondered whether these densely packed, presumably small cardiomyocytes enclosing residual scars were fully differentiated. While the embryonic form of myosin heavy chain, a marker for dedifferentiated cardiomyocytes (Sallin et al., 2015; Wu et al., 2016), was not detectable in the external wound border (Supplementary Figure 5B), cardiomyocytes in this region lacked assembled sarcomeres, and thus appeared to not be fully differentiated (Figure 4C).

**Figure 4.**
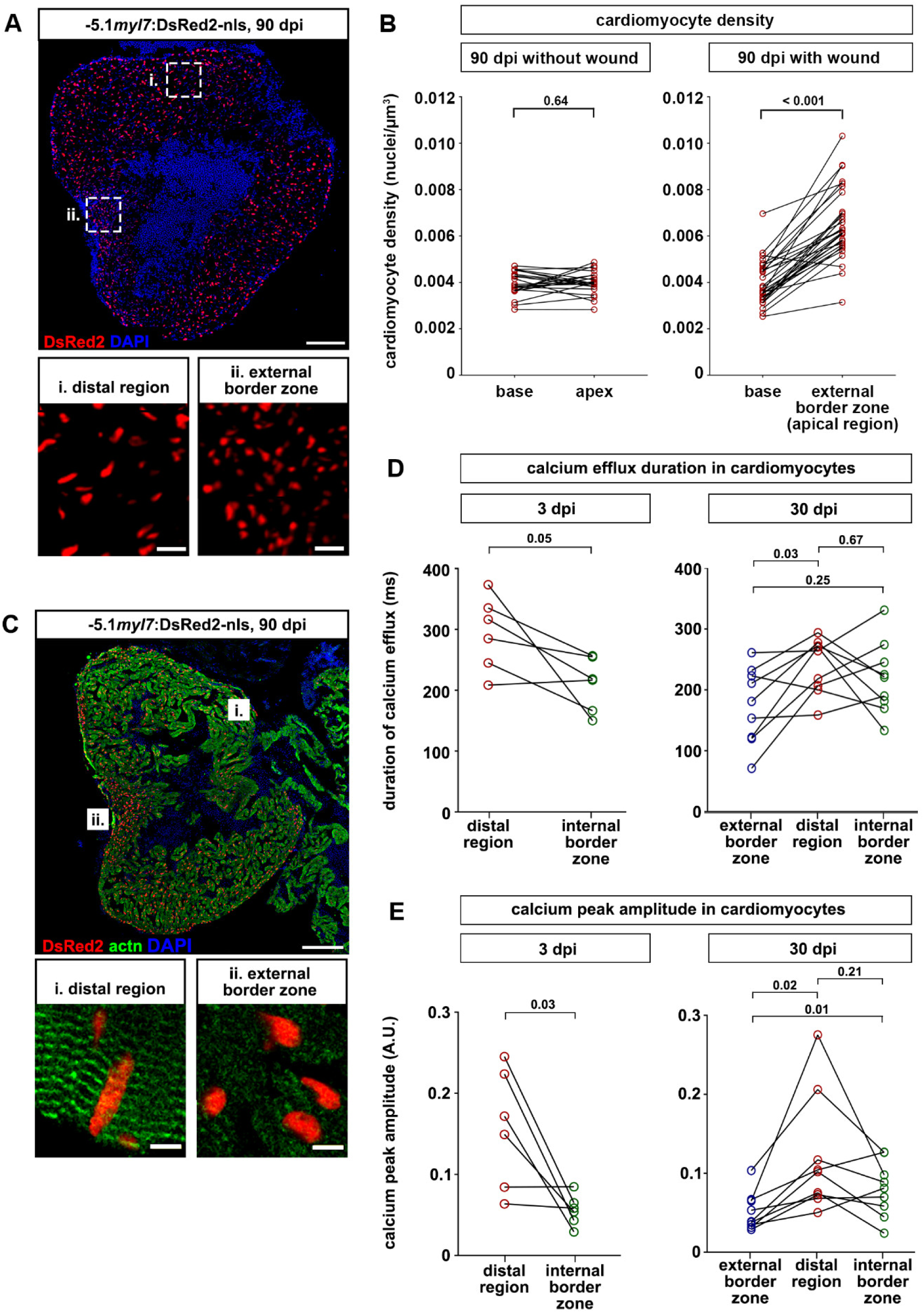
Cardiomyocytes are denser and less mature at the external wound border zone in scarred hearts. (A) Cardiomyocyte nuclei density is higher in the external wound border zone (box ii) than the distal region (box i) at 90 dpi. Scale bar, 100 µm. (B) At 90 dpi, cardiomyocyte density at base and apex does not differ in hearts without wound, but is higher at the external wound border in hearts with wounds. n (hearts) = 22 (without wound), 30 (wound). Paired two-tailed t-test. (C) Sarcomeric structures cannot be identified by alpha-actinin staining in the external wound border in hearts with residual scars. n (hearts with assembled sarcomeres): wound border, 2/13 hearts at 30 dpi, 4/23 hearts at 90 dpi; distal myocardium, 12/13 hearts at 30 dpi, 21/23 hearts at 90 dpi. Scale bar, 100 µm. (D, E) Calcium efflux duration (D) and amplitude of calcium peaks (E) in *myl7*:GCaMP6f-nls-T2A-RCaMP107-NES-pA)^hu11799^ cardiomyocytes is plotted at 3 dpi and at 30 dpi for hearts containing residual scars. n (hearts) = 6 (3 dpi), 9 (30 dpi). Paired two-tailed t-test.

Proper calcium handling can be used to evaluate the maturity of cardiomyocytes (Ronaldson-Bouchard et al., 2018). Calcium transients enter and leave cardiomyocytes in a cyclic fashion, resulting in calcium influx and efflux waves (van Opbergen et al., 2018). To measure calcium wave durations and amplitudes in regenerated hearts, we created transgenic fish expressing the calcium sensor GCaMP6f in cardiomyocytes, and established a system for live imaging of calcium transients on vibratome sections of adult hearts *ex vivo*. At 3 and 30 dpi, the duration of the calcium influx wave was not different at the internal or external wound border zone compared to the distal myocardial region (Supplementary Figure 5C). In contrast, at 3 dpi we observed a shorter duration of calcium efflux transients and a lower amplitude of calcium peaks at the internal wound border zone compared to the distal region, suggesting impaired functionality of dedifferentiating cardiomyocytes (Figure 4D, E). At 30 dpi, calcium efflux duration and peak amplitude of cardiomyocytes at the internal wound border zone were comparable to those in the distal region, indicating that these cells had re-differentiated and were as functional as cardiomyocytes in the region spared from injury (Figure 4D, E). However, in the external wound border zone calcium efflux duration was shorter and the amplitude of calcium peaks lower than in the distal region (Figure 4D, E). Together, these data suggest that regenerated cardiomyocytes at the external border of residual scars fail to fully mature.

Overall, our data show that restoration of cardiomyocyte numbers happens surprisingly fast within 30 days post cryoinjury and results in restoration of overall myocardial morphology. However, persistent fibrosis interferes with the differentiation of a subset of regenerated cardiomyocytes bordering residual scars (3-15% of the regenerated cardiomyocytes; 2,000 ± 800 at 30 dpi; 9,000 ± 1,000 at 90 dpi).

### Adult cardiomyocytes stay lineage-restricted during heart regeneration

Next, we wondered whether cardiomyocytes contribute to regeneration of other lineages in the heart. We generated a new transgenic Cre responder line *hsp70l*:LOXP-luc-myc-STOP-LOXP-dTomato, *cryaa*:YPet^ulm9^ (referred to hereafter as *hsp*:L to T), which expresses dTomato after excision of a *loxP*-flanked myc-tagged luciferase-Stop cassette following heat-shock (Figure 5A). Luc-myc was robustly expressed in all major lineages of cryoinjured hearts, namely cardiomyocytes (Figure 5C), epicardial cells (Figure 5D), endothelium (Figure 5E), neutrophils and macrophages (Figure 5F), and resident fibroblasts (Figure 5G, Supplementary Figure 6A) as well as activated fibroblasts (Figure 5H, Supplementary Figure 6A). Within each lineage, 30% to 60% of the cells expressed the transgene, based on measurement of the area positive for the lineage marker and luc-myc (Supplementary Figure 6B) and taking into account that the area positive for background staining of luc-myc was 10-15% in negative control fish (Supplementary Figure 6C). The absolute number of cells of a certain lineage expressing the transgene varied from 100 to 750 cells per heart section, in accordance with the different abundance of each cell type at 7 dpi (Supplementary Figure 6D, E). Based on these numbers we estimate that we should have been able to detect trans-differentiation rates of cardiomyocytes into other lineages as low as 0.3 - 2%, depending on the cell type. Importantly, within each lineage the cells expressing the *hsp*:L to T transgene did not appear to be clustered in a spatially restricted subset of cells (Supplementary Figure 7). A statistical approach that assesses signal correspondence between two numerically quantified images indeed showed concordance between areas expressing the lineage marker GFP and areas expressing both GFP and luc-myc for lines marking the endocardium, epicardium and macrophages and neutrophils (Supplementary Figure 8 and Supplementary Table 1). We conclude that the transgene is not expressed in a specific subset of cells within each lineage, making it unlikely that our tracing experiments would have missed transdifferentiation into any potentially existing sub-lineages. While statistical analysis did not support concordance of luc-myc expression with the *wt1b*:EGFP^li1Tg^ line, which labels mainly epicardium and fibroblasts (Sanchez-Iranzo et al., 2018) (Supplementary Table 1), and was not possible for the other fibroblasts markers due to low cell counts, the fact that our tracer *hsp*:L to T transgene was co-expressed with 5 different markers of fibroblasts (Figure 5G, H, Supplementary Figure 6A) makes it unlikely that we would have missed transdifferentiation into fibroblasts.

**Figure 5.**
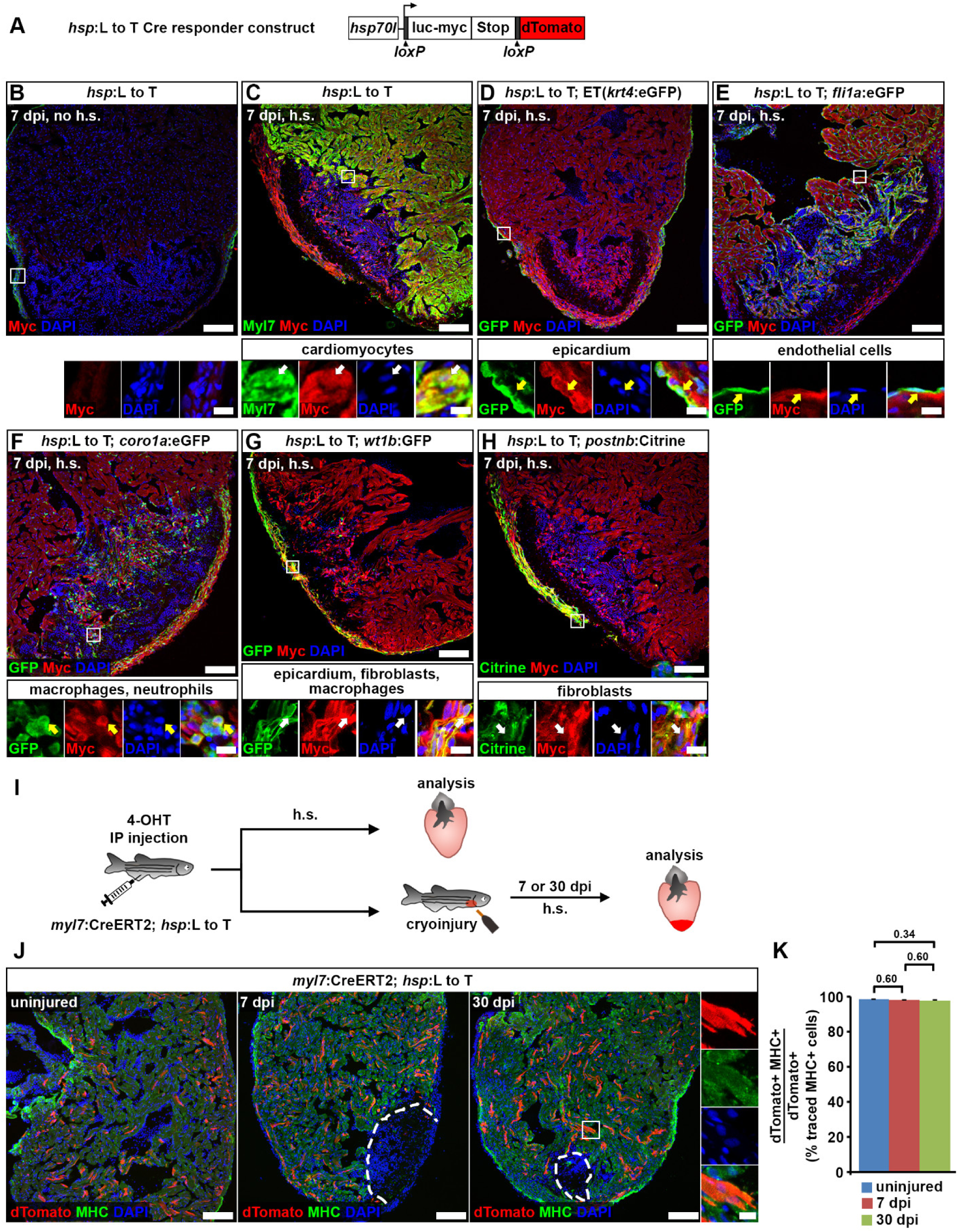
Cardiomyocytes remain lineage-restricted during adult heart regeneration. (A) Schematic of the *hsp70l:*LOXP-luc-myc-STOP-LOXP-dTomato, *cryaa:*YPet^ulm9^ (*hsp*:L to T) transgene. (B-H) At 7 dpi, myc-tagged luciferase is not detectable in the ventricle of adult *hsp70l*:LOXP-luc-myc-STOP-LOXP-dTomato, *cryaa*:YPet^ulm9^ (*hsp*:L to T) hearts without heat-shocks (B). After heat-shock, expression is found in *myl7*^+^ cardiomyocytes (C), Et(*krt4*:eGFP^sqet27^)^+^ epicardial cells (D), *fli1a*:eGFP^y1Tg +^ endocardial cells (E), *coro1a*:eGFP^hkz04tTg +^ leukocytes (F), in *wt1b*:EGFP^li1Tg +^ epicardial cells, fibroblasts and macrophages (G), and in *postnb*:Citrine^cn6Tg +^ fibroblasts (H). Boxed areas are shown in magnified view. Scale bars, 100 µm (overview), 10 µm (magnified view). (I) Experimental design for adult cardiomyocyte lineage tracing. (J) dTomato^+^ traced cells are randomly located throughout the ventricle, and express the cardiomyocyte marker myosin heavy chain (MHC). Scale bars, 100 µm (overview), 12.5 µm (magnified view). (K) No difference is detected in average percentages of dTomato^+^ traced cells expressing MHC between uninjured and regenerating hearts. Error bars, s.e.m. n (hearts) = 4 (uninjured), 4 (7 dpi), 5 (30 dpi). n (traced cells) = 942 (uninjured), 775 (7 dpi), 1134 (30 dpi). One-way ANOVA + Holm-Sidak’s multiple comparisons test. Observed difference uninj. vs 7 dpi = 0.3 (0.3% of traced cells in uninj.); smallest significant detectable difference = 0.7 (0.7% of uninj). Observed difference uninj. vs 30 dpi = 0.6 (0.6% of traced cells in uninj.); smallest significant detectable difference = 1.2 (1.2% of uninj.). Observed difference 7 dpi vs 30 dpi = 0.2 (0.2% of traced cells in 7 dpi); smallest significant detectable difference = 1.3 (1.4 of 7 dpi).

We labelled cardiomyocytes in adult fish prior to heart injury using *cryaa*:DsRed,*-*5.1*myl7*:Cre-ERT2^pd10^ (*myl7*:CreERT2) (Kikuchi et al., 2010) together with *hsp*:L to T (Figure 5I). dTomato^+^ cells were randomly distributed throughout the entire ventricle, indicating that recombination resulted in labelling of a random subset of cardiomyocytes (Figure 5J, K). In regenerating hearts at 7 and 30 dpi, 98% of all dTomato^+^ cells were myosin heavy chain (MHC)^+^ cardiomyocytes (Figure 5J, K). While a small fraction of traced cells could not unequivocally be identified as cardiomyocytes (1.5 ± 0.1% in uninjured hearts), this fraction did not differ between uninjured and injured hearts (Figure 5K), and thus most likely reflected technical limitations of registering co-staining of MHC and dTomato rather than transdifferentiation. Thus, we conclude that adult cardiomyocytes do not significantly contribute to other cellular lineages during regeneration.

### Cardiomyocytes do not transdifferentiate in juvenile regenerating hearts

The cellular mechanisms of regeneration can change with age (Chera et al., 2014). Thus, we assessed ventricular cardiomyocyte plasticity in juveniles. We found that cryoinjured hearts of juvenile fish (standard length ∼ 11 - 16 mm (Parichy et al., 2009)) robustly resolved wound tissue within 30 dpi (Figure 6A, B). Juvenile cardiomyocyte cell cycle activity peaked at 7 dpi and was hardly detectable by 30 dpi, similar to adult hearts (Figure 6C, D).

**Figure 6.**
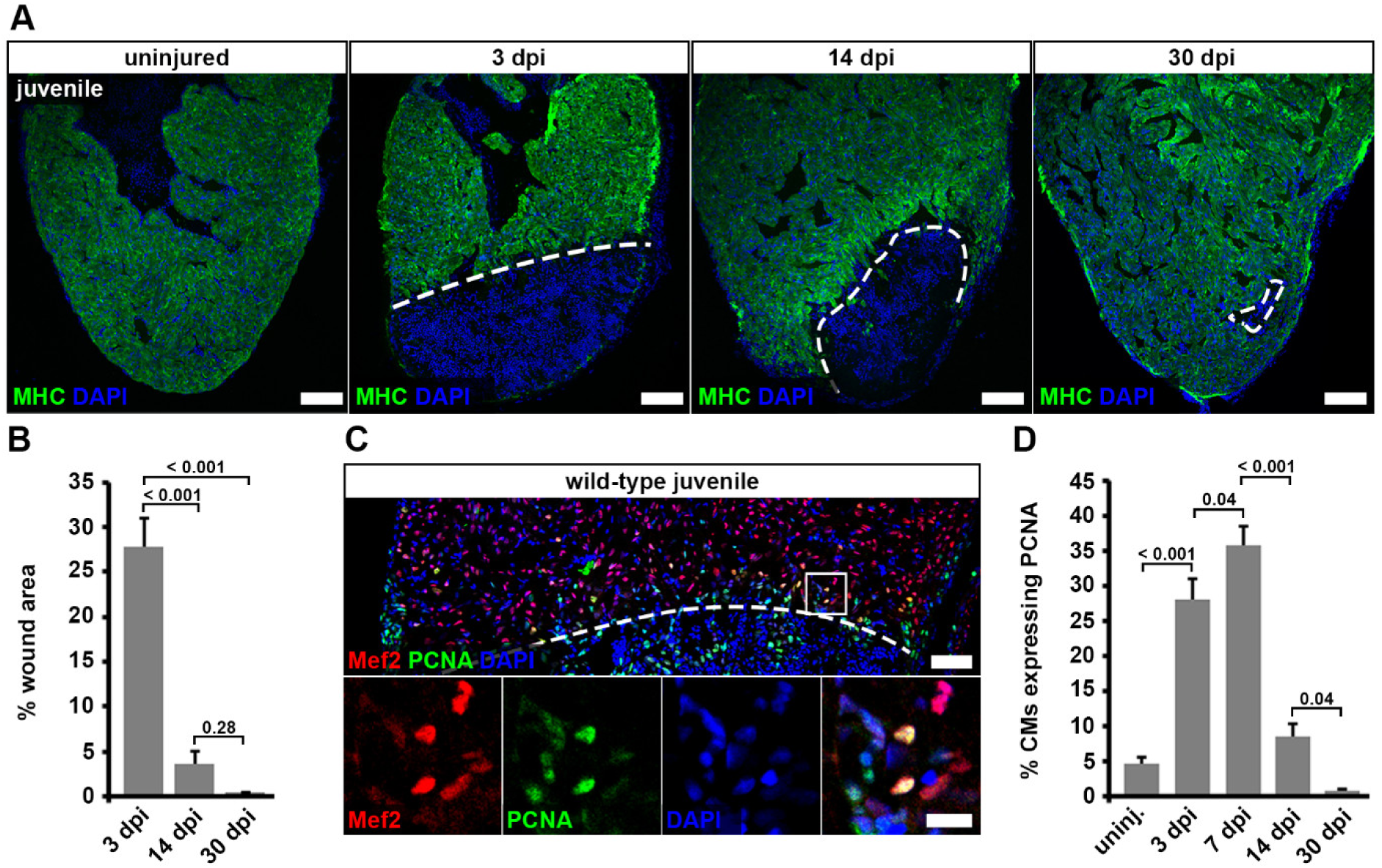
Juvenile hearts regenerate after cryoinjury via cardiomyocyte proliferation. (A) Myosin heavy chain (MHC) immunostaining reveals wound area (dashed lines) on heart sections of cryoinjured juvenile fish. Scale bars, 100 µm. (B) Wound area (MHC^—^ area) relative to the whole ventricle area is plotted. Error bars represent s.e.m. n (hearts) = 6 (3 dpi), 6 (14 dpi), 6 (30 dpi). One-way ANOVA + Holm-Sidak’s multiple comparisons test. (C) PCNA^+^ Mef2^+^ cardiomyocytes in juvenile hearts at 7 dpi. Scale bars, 50 µm (overview), 12.5 µm (magnified view). (D) Average percentage of cardiomyocytes expressing PCNA within 150 µm from the wound border is plotted. Error bars represent s.e.m. n (hearts) = 6 (uninjured), 6 (3 dpi), 6 (7 dpi), 6 (14 dpi), 6 (30 dpi). One-way ANOVA + Holm-Sidak’s multiple comparisons test.

We combined *myl7*:CreERT2^pd10^ with another tracer line, *hsp70l*:LOXP-DsRed2-LOXP-NLS-EGFP^tud9^ (*hsp*:R to nG), which expresses nuclear GFP after excision of a DsRed2-stop cassette (Knopf et al., 2011). DsRed could be detected in endothelial and epicardial cells, blood cells, cardiomyocytes, and resident and activated fibroblasts (Figure 7A-D and Supplementary Figure 9A). Thus, this tracer line also allowed us to test whether cardiomyocytes give rise to other cell types during regeneration. Recombination was induced in embryos and fish were raised to juvenile stages (Figure 7E). This protocol resulted in labeling of 70% of cardiomyocytes (Supplementary Figure 9B). In uninjured hearts, 99.4 ± 0.4% of all analyzed GFP^+^ cells expressed Mef2 and were hence cardiomyocytes (Figure 7F, G). Similarly, 99% of GFP^+^ traced cells were Mef2^+^ at 7 and 30 dpi (Figure 7G). Thus, we conclude that both adult and juvenile cardiomyocytes remain lineage-restricted during zebrafish heart regeneration.

**Figure 7.**
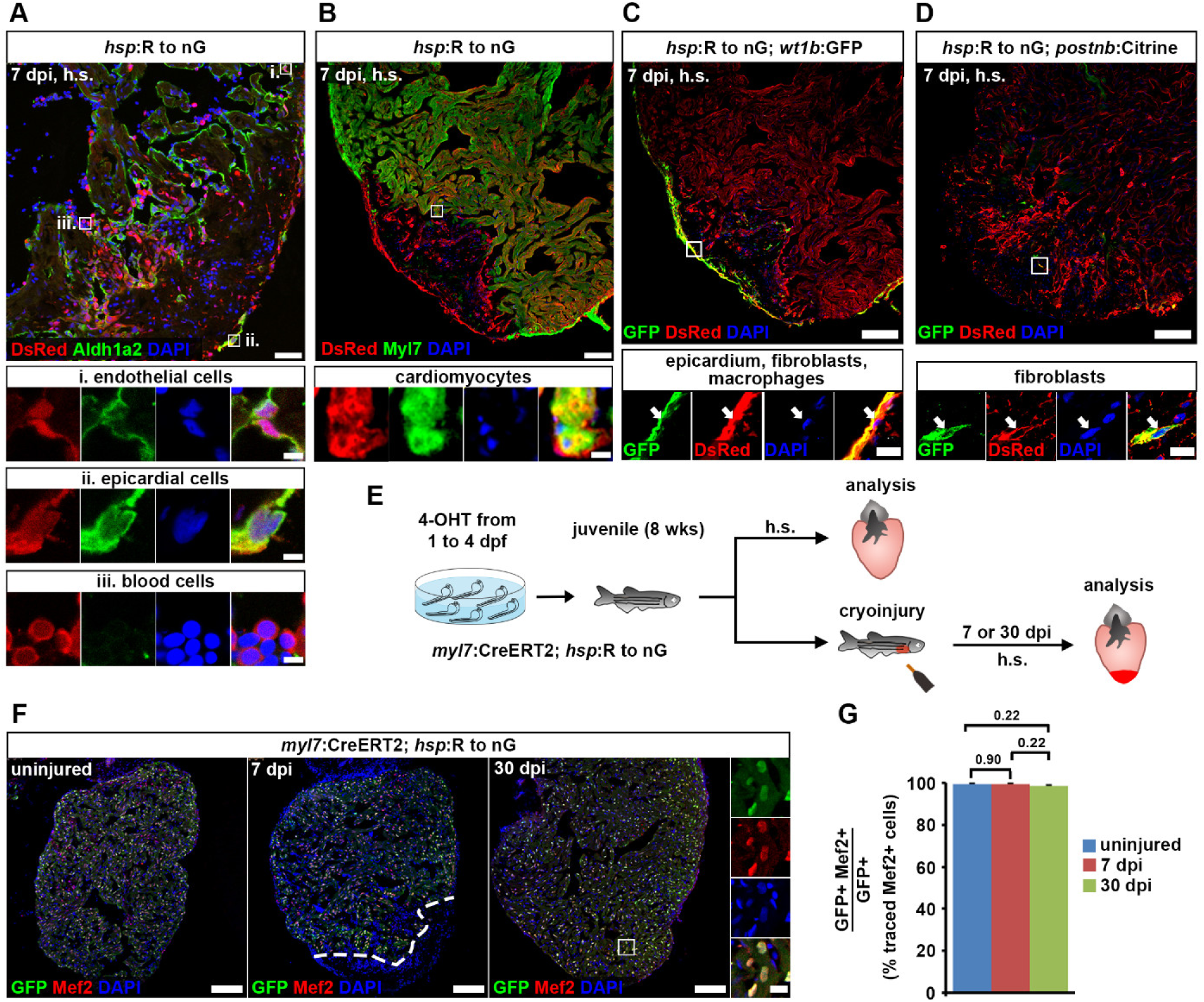
Cardiomyocytes remain lineage-restricted during juvenile heart regeneration. (A-D) At 7 dpi in juvenile fish, expression of the *hsp*:R to nG^tud9^ tracer transgene is detected by DsRed immunofluorescence after heat-shock (h.s.) in endocardial cells (Aldh1a2^+^, boxed area i in (A)), epicardial cells (Aldh1a2^+^, boxed area ii in (A)), blood cells (Aldh1a2^-^ with small, round and intense DAPI-stained nuclei, boxed area iii in (A)), cardiomyocytes (Myl7^+^, boxed area in (B)), *wt1b*:EGFP^li1Tg +^ epicardial cells, fibroblasts and macrophages (C), and in *postnb*:Citrine^cn6Tg +^ fibroblasts (D). Scale bars, 40 µm (overview), 5 µm (magnified view). (E) Experimental design for juvenile cardiomyocyte lineage tracing. (F) Nuclear GFP^+^ traced cells are widespread throughout the ventricle and colocalize with Mef2, a nuclear cardiomyocyte marker. Scale bars, 100 µm (overview), 12.5 µm (magnified view). (G) No significant difference is detected in average percentages of GFP^+^ cells expressing Mef2 between uninjured and regenerating hearts. Error bars, s.e.m. n (hearts) = 4 (uninjured), 5 (7 dpi), 6 (30 dpi). n (traced cells) = 858 (uninjured), 823 (7 dpi), 1123 (30 dpi). One-way ANOVA + Holm-Sidak’s multiple comparisons test. Observed difference uninj. vs 7 dpi = 0.06 (0.06% of traced cells in uninj.); smallest significant detectable difference = 0.8 (0.8% of uninj). Observed difference uninj. vs 30 dpi = 0.8 (0.8% of traced cells in uninj.); smallest significant detectable difference = 1.9 (1.9% of uninj). Observed difference 7 dpi vs 30 dpi = 0.9 (0.9% of traced cells in 7 dpi); smallest significant detectable difference = 1.6 (1.6% of 7 dpi).

## Discussion

### A model for complete heart regeneration by proliferation of lineage-restricted cardiomyocytes

Based on our results and previous findings we propose the following model for zebrafish heart regeneration (Figure 8). In response to heart injuries, spared differentiated cardiomyocytes at the wound border dedifferentiate and re-enter the cell cycle (Chablais and Jazwinska, 2012; Gonzalez-Rosa et al., 2011; Kikuchi et al., 2010; Poss et al., 2002; Schnabel et al., 2011; Wang et al., 2011; Wu et al., 2016). Dedifferentiation entails the loss of features of the mature state like sarcomeric structures, alterations in calcium handling, and the gain of markers associated with cardiogenesis during development, but does not include acquisition of multipotency (Figure 8A). In response to cryoinjuries that kill about 1/3 of the cardiomyocytes, restoration of pre-injury cardiomyocyte numbers is already completed by 30 days post injury, and results in regeneration of myocardial morphology, in particular the septation of trabecular and compact myocardial layers (Figure 8B). While cardiomyocyte numbers are very robustly regenerated, full scar resolution is only observed in 30% of the injured hearts at 30 or 90 dpi. Persistent fibrosis leads to improper differentiation of a subset of regenerated cardiomyocytes located at the external wound border (Figure 8B).

**Figure 8.**
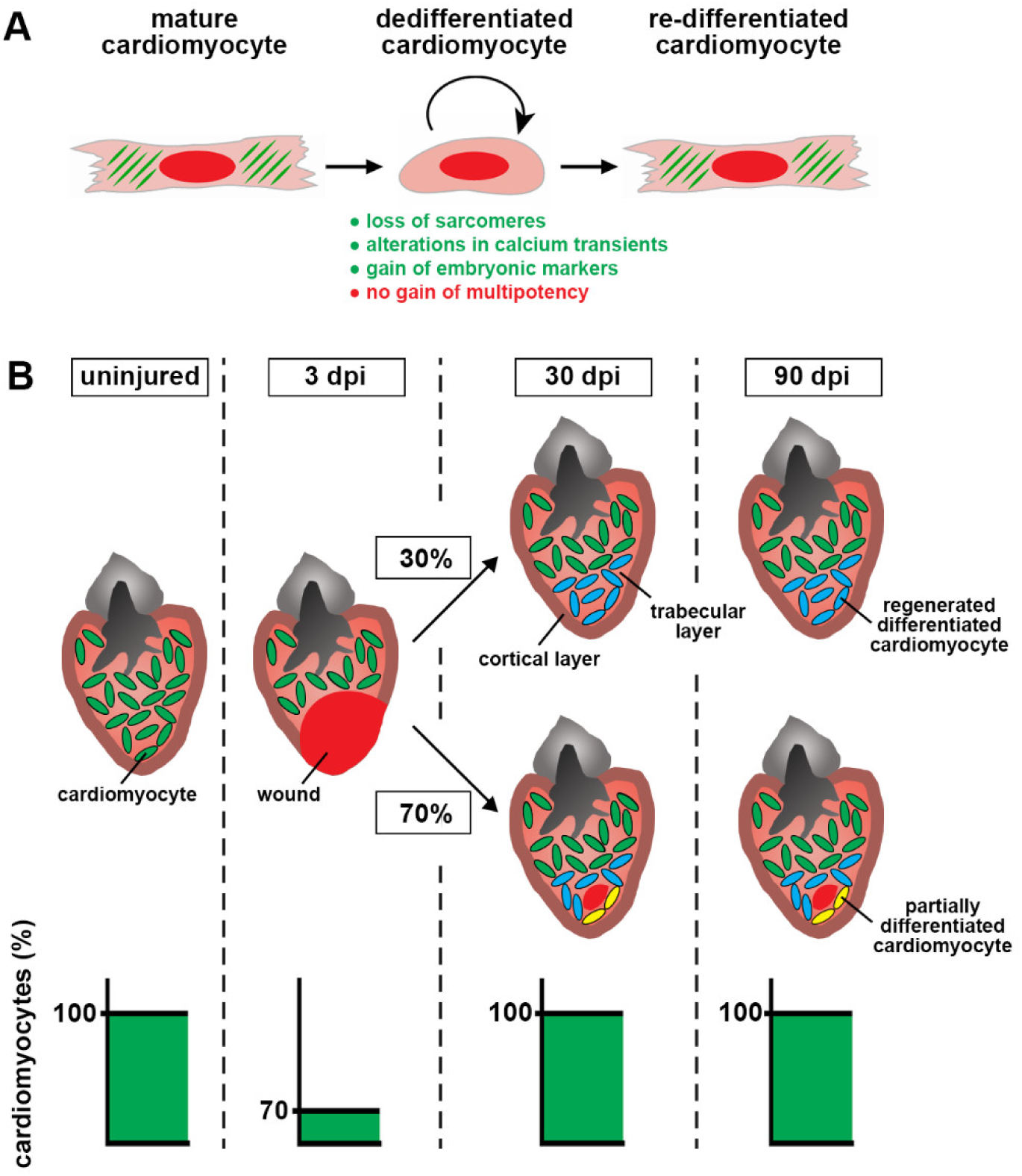
Quantitative model of zebrafish heart regeneration via proliferation of lineage-restricted cardiomyocytes. (A) Model of cardiomyocyte fate during zebrafish heart regeneration. Cardiomyocytes dedifferentiate, which entails loss of sarcomeres, alterations in calcium transients and gain of embryonic markers, but not attainment of multipotency. (B) Model of the time-courses of cardiomyocyte regeneration and wound resorption. Restoration of pre-injury cardiomyocyte numbers (green) in response to cryoinjury, which kills about 30% of the cardiomyocytes, is achieved within 30 days. The morphological septation between trabecular layer (pink) and cortical layer (brown) is restored. Despite completed cardiomyocyte regeneration, the majority of hearts still contain scars at 30 and 90 dpi. Regenerated cardiomyocytes in scar-free ventricles or at the internal wound border zone retain normal density and fully re-differentiate (cyan). Some of the regenerated cardiomyocytes (3-15%) enclosing the remaining internalized scars are smaller and only partially re-differentiated (yellow). In scarred hearts, fibrosis persists beyond 30 dpi.

### Scar resolution and cardiomyocyte regeneration

Our results suggest a more complex answer to the question whether zebrafish can “completely” regenerate the heart than previously appreciated. While regeneration of cardiomyocyte numbers is indeed complete when analyzed on the level of a population of fish, some defects in cardiomyocyte maturation and scar resolution remain in response to cryoinjuries in many hearts, which is consistent with the finding that ECM-producing fibroblasts are incompletely eliminated (Sanchez-Iranzo et al., 2018). One caveat in our study is the impossibility to unambiguously determine whether each individual heart with a residual wound has regenerated all cardiomyocytes. Despite this limitation, our findings indicate that cardiomyocyte proliferation is uncoupled from full scar resorption, since cardiomyocytes appear to withdraw from the cell cycle within 30 days post injury irrespective of whether the scar has fully resolved.

The limited temporal window of cardiomyocyte cell cycle activity after injury is consistent with the idea that all cardiomyocytes that show cell cycle activity indeed proliferate during regeneration. In fact, a simple mathematical model that is based on the assumptions that all PCNA^+^ cardiomyocytes that are present at the wound border at different time-points after injury proliferate, and that cell cycle length is 24 hours, predicts complete cardiomyocyte regeneration within 30 days post injury, which is in agreement with our experimental data (Supplementary Figure 10A). Thus, cardiomyocyte cell cycle activity (PCNA or S-phase markers) can be used as a proxy for the extent of cardiomyocyte regeneration. Of note, even at the peak of cardiomyocyte cell cycle activity (7 dpi), where on average ∼100 PCNA^+^ cardiomyocytes can be detected per heart section, only very few cardiomyocytes (0-2 per section) stain positive for phospho-Histone 3. This indicates that mitosis - or at least the period in which H3 is phosphorylated - takes less than half an hour.

Increased cardiomyocyte density in the external wound border has been reported before, and was interpreted as evidence of myocardial hyperplasia (Gonzalez-Rosa et al., 2014). Our study confirms a local increase in cardiomyocyte density around the residual wound, and in addition shows that these cardiomyocytes are not fully mature, since they lack organized sarcomeric structures and show defects in calcium signaling. In our hands, these cardiomyocytes appear to be located in the trabeculated myocardium, and we find no evidence for a thickening of the compact layer of the myocardium, in contrast to previous reports (Gonzalez-Rosa et al., 2014; Sanchez-Iranzo et al., 2018). While we have used a specific molecular marker for the cortical layer, these other studies have relied on histology, which might limit the ability to distinguish trabecular and cortical layers. Since the thickness of the cortical layer remains constant after regeneration, our data imply that dense cardiomyocytes in the regenerated ventricular wall are smaller (Figure 8B). While the localized regions of partially differentiated cardiomyocytes account for only 1-5% of the total cardiomyocyte number (3-15% of regenerated cardiomyocytes), their presence might explain why several studies have found that some aspects of heart function do not fully recover, in particular movement of the ventricular wall remains altered in cryoinjured hearts (Gonzalez-Rosa et al., 2014; Gonzalez-Rosa et al., 2011; Hein et al., 2015; Wang et al., 2011).

Although we detect immature cardiomyocytes and residuals scars in many regenerated zebrafish hearts, these defects are small, with ∼85-97% of regenerating cardiomyocytes attaining full differentiation and ∼90% of wound tissue being fully resorbed. Together with the fact that cardiomyocyte numbers completely regenerate at the population level, our results confirm the high regenerative capacity of the adult zebrafish heart, which stands in stark contrast to the adult mammalian and even neonatal mouse heart, where cryoinjuries similar to those performed in zebrafish do not result in any regenerative responses (Darehzereshki et al., 2015; Polizzotti et al., 2016; Polizzotti et al., 2015).

### Cardiomyocytes are lineage-restricted during zebrafish heart regeneration

Several previous studies have established that differentiated cardiomyocytes dedifferentiate and represent the source of regenerating cardiomyocytes in the zebrafish heart (Gupta et al., 2013; Gupta and Poss, 2012; Jopling et al., 2010; Kikuchi et al., 2010). Our work now indicates that cardiomyocytes do not dedifferentiate to a multipotent state, but rather remain lineage-restricted and only form cardiomyocytes during heart regeneration. We could not detect cardiomyocyte markers in 1.5 ± 0.1% of traced cells in uninjured adult hearts and in 0.6 ± 0.4% of cells in uninjured juvenile hearts, which most likely is due to failure to detect existing co-expression, caused by immunostaining and/or imaging issues. We estimate that we should have been able to unambiguously detect transdifferentiation above this background if it had occurred in at least 5% of adult, and 2.5% of juvenile cardiomyocytes. The recombination efficacy that we could achieve in adult fish was modest, thus it is possible that we failed to trace a rare subpopulation of cardiomyocytes that would be able to transdifferentiate. However, we consider it unlikely that such a subpopulation exists, since we also could not detect transdifferentiation in juvenile fish, where recombination efficiencies exceeded 70% after tamoxifen treatments in embryos. Secondly, both in adult and juvenile fish, all analyzed cell types expressed our tracer transgenic lines and recombined cardiomyocytes were randomly distributed within the myocardium, making it unlikely that we missed a cardiomyocyte subpopulation located in a specific anatomical region. Based on the number of traced cardiomyocytes that we could not unequivocally co-localize with cardiomyocyte markers, the number of cells of the individual non-myocyte cellular lineages present in each heart, and the fraction of these cells that express our tracer transgenes, we estimate that in adult fish we should have been able to detect transdifferentiation rates as follows, where the numbers represent the minimum fraction of non-myocytes originating from traced cardiomyocytes: 0.7% for *krt4*^+^ epicardial cells, 0.3% for *fli1a*^+^ endothelial cells, 0.4% for *coro1a*^+^ leukocytes, 1.3% for *wt1b*^+^ epicardial cells, fibroblasts and macrophages, and 2% for *postnb*^+^ fibroblasts. In juveniles, we suppose that such percentages should have been even lower, given the highly ubiquitous expression of the tracer in the entire ventricle and the lower uncertainty than in adults in registering traced cells as cardiomyocytes. We conclude that cardiomyocyte transdifferentiation does not occur or is a very rare event during unperturbed heart regeneration.

Our results thus add to the growing body of evidence that attainment of multipotency is a rare cellular event during unperturbed vertebrate regeneration. Thus, efficient heart regeneration does not require such specialized lineage reversal, raising the hope that heart regeneration can be therapeutically induced in humans.

## Acknowledgements

We thank Christa Haase and Doris Weber for creation of the *hsp*:L to T transgene, and Doris Weber, Brigitte Korte and Martina Raasholm for fish care and technical help.

