## Supplemental Methods, Figures and Table for "Is zebrafish heart regeneration “complete”? Lineage-restricted cardiomyocytes proliferate to pre-injury numbers but some fail to differentiate in fibrotic hearts"

### Bertozzi et al, Supplemental Materials

**Contents**


| | |
|---|---|
| Expanded Materials & Methods | page 2 |
| Supplementary Figures and Figure legends | page 6 |
| Supplementary Table | page 21 |

### Expanded Materials & Methods

*Cell number quantifications*

Quantifications for fate mapping and cardiomyocyte proliferation experiments were performed manually with ImageJ software. Traced cardiomyocytes were analyzed on 4 to 7 sections per heart. In adult fate mapping experiments using the *hsp70l*:LOXP-luc-myc-STOP-LOXP-dTomato, *cryaa*:YPet$^{ulm9}$ (*hsp*:L to T) responder line, each nucleus that was surrounded by unequivocally co-localized dTomato and MHC on several optical z-planes was counted as a traced cardiomyocyte. In juvenile fate mapping experiments using the *hsp70l*:LOXP-DsRed2-LOXP-NLS-EGFP$^{tud9}$ (*hsp*:R to nG) responder line, cells displaying nuclear co-localization of GFP with Mef2 on single optical plane images were counted as traced cardiomyocytes. The fact that the latter nuclear co-localization is easier to detect unequivocally than the cytoplasmic co-localization in the *hsp*:L to T responder likely explains why we counted a slightly higher percentage of recombined cells to be of cardiomyocyte character in the juvenile vs adult experiments (99% vs 98%). PCNA expression in cardiomyocytes was analyzed on 2 to 3 sections per heart displaying the biggest wounds. Proliferation indices were calculated either from cardiomyocytes situated within 150 µm from the wound border in injured samples or from all cardiomyocytes in uninjured samples.

For cardiomyocyte quantification experiments, all sections on 1/6$^{th}$ of the slides, over which the sections of each heart were distributed, were analyzed. If heart sections were distributed on 12 or 18 slides, 2 or 3 non-consecutive slides were selected, respectively. Image analysis was performed with ImageJ. Automated quantification of cardiomyocyte number was performed using the Find Maxima function. Noise tolerance was set to a value which creates a single point selection for each cardiomyocyte nucleus based on native DsRed2 fluorescence, while giving no false positive selections in the background. Imaging conditions and noise tolerance parameters were kept constant within the same experimental pool, as well as among the three main experiments with different final time-points. Estimation of the total number of cardiomyocytes in the whole ventricle was performed by multiplying the measured number of cardiomyocytes by 6 (since we analyzed 1/6th of sections) and by multiplying with 0.715, a factor designed to correct for the fact that a certain number of cardiomyocyte nuclei are shared between adjacent sections, since they were cut through. To determine this correction factor, we produced 50 µm thick cryosections, acquired confocal stacks of the entire section and counted the number of nuclei within a 10 µm-thick sub-stack. Cardiomyocyte nuclei present in the first and last optical plane of the sub-stack were considered severed if they were detected also in the adjacent optical planes of the original 50 µm stack. This analysis showed that 57% of the cardiomyocyte nuclei are cut through and thus shared between sections, while 43% are not shared between sections. We then decided to assign all shared cardiomyocytes in the bottom optical plane (half of the cut cardiomyocytes, i.e. 28.5%) to the same analyzed sub-stack, while all shared cardiomyocytes in the top optical plane (the other 28.5%) were assigned to the optical planes above the sub-stack. Thus only 43% plus 28.5% (71.5%) of all counted cardiomyocytes were considered unique to a 10 µm-thick sub-stack, and therefore the same ratio was applied to correct the number of cardiomyocytes in each physical 10 µm section. For comparison of manual and automatic counting and for analysis of superimposed cardiomyocyte nuclei, a 150x150 µm region was randomly selected in 1-2 sections from 3-4 hearts. Manual and automatic counting of confocal images was performed on maximum projections of z-stacks. Superimposed cardiomyocytes were quantified by analyzing overlapping nuclei in two focal planes 5 µm apart of z-stack confocal images.

*Ventricular area, wound area and ventricular volume quantification*

Total size of the ventricular area and wound area was measured by adding up the individual areas of all sections on one or more non-consecutive serial slides representing 1/6$^{th}$ of the ventricle. For juvenile hearts, wound area was defined as the MHC$^-$ region of the ventricle, and ventricular area was defined as the whole MHC$^+$ and MHC$^-$ area combined. For cardiomyocyte quantification experiments, wound area was defined as the DsRed2$^-$ region of the ventricle, and ventricular area was defined as the whole DsRed2$^+$ and DsRed2$^-$ area combined. Total ventricle volume was calculated by multiplying the measured ventricular area by 6*10 µm.

*Quantification of cardiomyocyte density and compact layer width*

Cardiomyocyte density was calculated by dividing the number of cardiomyocyte nuclei by the volume containing the selected cardiomyocytes. The apex volume was defined as the entire region containing the myocardial layer that forms the external wall surrounding the remaining wound, while the base volume was defined as a ~ 250 µm x 250 µm region most distal from the apex, in close proximity to the bulbus arteriosus. Width of the myocardial compact layer was calculated by manually outlining the region of the myocardium labeled with laminin with an inner and outer line, and averaging the minimum distance between all coordinate points of the two lines. Width was measured for the compact layer surrounding the apex of the ventricle or only for the portion encompassing the wound edge, defined as the region of the regenerated myocardium which encloses any internalized wound of hearts with a remaining scar.

*Modeling of cardiomyocyte regeneration time-course*

Theoretical numbers of regenerated cardiomyocytes were calculated based on the PCNA timecourse (Figure 1A), according to the following equation:

$$\%\,CM(n) = \%\,CM(0) * [\prod_{t=0}^{n}(1+F_t)^t]^{(1/i)}$$

where $n$ is the time point (days post injury), $t$ is the time interval (days) between two consecutive time points of the PCNA timecourse, $F_t$ is the cardiomyocyte proliferation rate during $t$ and $i$ the cell-cycle length in days. $F_t$ values were calculated by taking all the combinations of two consecutive time points of the PCNA timecourse and averaging the PCNA$^+$ cardiomyocyte rates of those time points. Therefore, only 5 values for $F_t$ (calculated from the 5 time intervals between consecutive time points) were used to estimate the number of new cardiomyocytes generated at any given day post injury. Values were calculated assuming that all PCNA$^+$ cardiomyocytes divide and assuming a constant cell-cycle length of 24 hours ($i$=1). Curve fitting was performed using the built-in exponential recovery function of ImageJ.

*Sensitivity of detection for trans-differentiation rates*

The average number of traced Tomato$^+$ cells that could not be unambiguously identified as cardiomyocytes based on MHC staining was measured as 1 cell per section in uninjured adult hearts, which constitutes the background limiting the sensitivity to detect cardiomyocyte trans-differentiation during regeneration. From this mean, SD of values and sample size, we computed the minimum effect size that could be significantly detected in the adult lineage tracing experiments. Such effect size corresponded to a difference of 1 cell per section above the background, which translates into a minimum of 2 traced cardiomyocytes per section required to detect a single trans-differentiation event. Thus, the lowest measurable trans-differentiation rate equals to 2 myocytes divided by the number of non-myocytes. Such rate corresponds to 0.3 – 2%, given that we could image between 750 and 100, respectively, non-myocyte cells per section, depending on the abundance of each cell type at 7dpi.

*Analysis of cardiomyocyte calcium cycling*

At the selected time-points, cryoinjured *myl7*:GCaMP6f-nls-T2A-RCaMP107-NES-pA$^{hu11799}$ transgenic hearts were extracted and placed in calcium loading buffer (CLB, 125 mM NaCl, 20 mM HEPES pH 7.4, 10 mM glucose, 4.5 mM KCl, 2 mM $MgCl_2$, and 1.5 mM $CaCl_2$, pH 7.4) containing 10 mM 2,3-butanedione monoxime (BDM) in order to uncouple the excitation-contraction process. Ventricles were mounted in a plastic cryomold containing 1% agarose in PBS and 10 mM BDM. Agarose blocks were sectioned on a vibratome filled with room temperature PBS with 10 mM BDM at 150-200 µm thickness. The settings on the vibratome were: frequency = 80, amplitude = 1.0, velocity = 10. Slices were transferred into fresh CLB containing 10% fetal calf serum (FBS) and 10 mM BDM. The agarose surrounding the slices was carefully removed and placed on a glass coverslip that had been coated with poly-L-lysine. A drop of 0.5% agarose in CLB with 10 mM BDM was placed on top of the slice to hold it in place. Mounted slices were placed in a recording chamber (28 ºC) containing CLB with 10% FBS and 10 mM BDM. Platinum electrodes were place nearby the slice and electrical stimulation was performed using a Stimulator CS (Hugo-Sachs; Harvard Apparatus) set at 1 Hz. Slices were imaged on a custom-built upright epifluorescence microscope with a 20X, 1.0 NA Olympus dipping objective and a high speed fluorescence-detecting camera. A custom micromanager software (ImageJ) was used to record experiments and files were saved as a tiff series. For analysis, a 10x10 µm selection window in Fiji was used to measure fluorescence fluctuations over time and saved as a csv file. Two regions per area of interest were used as technical replicates. Custom matlab scripts were used to measure the dynamics of the calcium wave. Calcium influx was defined as the time required to go from 10 to 90% of maximal calcium transient, while efflux was defined as the time needed to go from 90% to 10% of the calcium transient. Peak amplitude was defined as the relative amount of calcium transients from Resting Membrane Potential (RMP) to its Maximum peak (Max).

*Statistical Concordance Analysis*

The entire analysis was performed in the R statistical computing environment (R Core Team, 2019). The images were first imported into R using the package imager (Barthelme, 2020) and the quantitative matrix was binary coded, where 1 indicates a signal and 0 represents absence of signal. To facilitate downstream quantitative analysis by reducing image size (Peltier, 2018), we binned individual binary pixel counts by taking the square root of the total number of pixel count in a given image. Based on the number of binary positive counts in a given bin, regions of low, intermediate, and high were identified using Ward's method, a clustering approach that minimizes the total within-cluster variance and forms distinct groups (Ward, 1963; Murtagh and Legendre, 2014). Concordance/similarity between GFP and myc high pixel count regions was performed using Kendall's coefficient of concordance (Kendall and Babington Smith, 1939; Kendall and Gibbons, 1990). The statistical significance is based on the calculated Chi statistic and the level of concordance is measured by Kendall's coefficient, ranging between 0 and 1 (1 represents complete concordance).

### Supplementary Figures and Figure legends

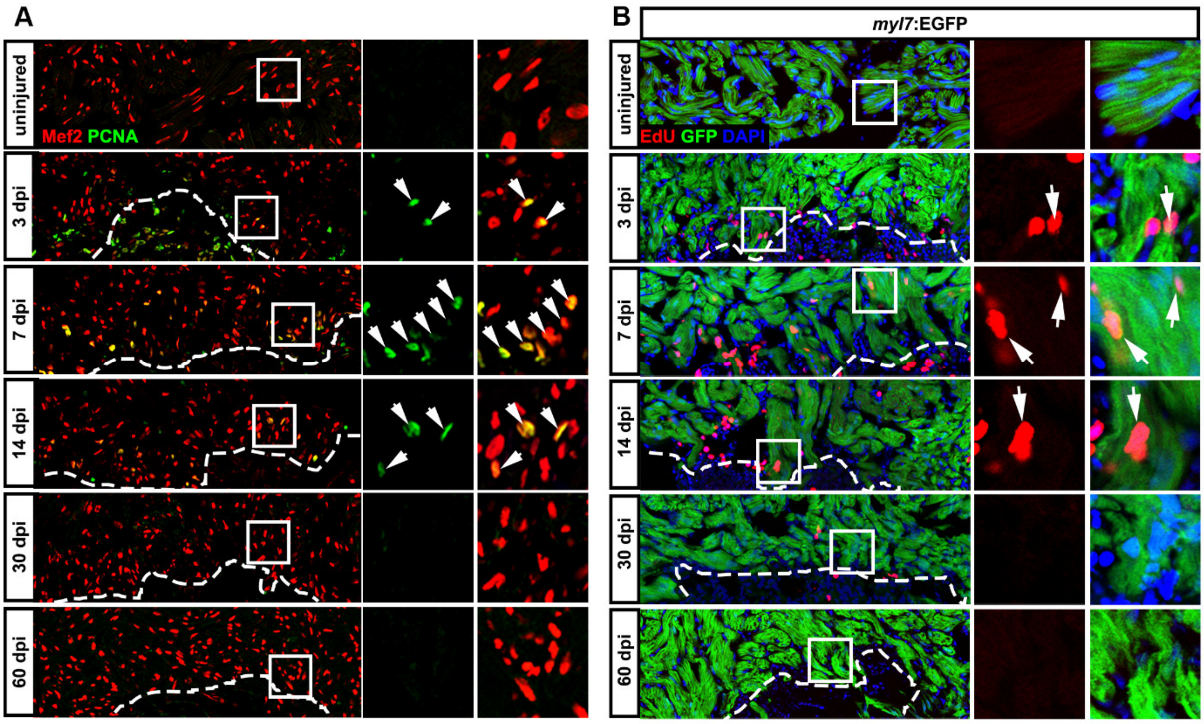


**Supplementary Figure 1. The fraction of PCNA$^{+}$ and EdU-incorporating cardiomyocytes reaches baseline by 30 dpi.**

(A) Representative images of PCNA immunofluorescence revealing cycling Mef2$^{+}$ cardiomyocytes (arrowheads) at the wound border (dashed line) at different time points after cryoinjury. Quantification of these data is found in Figure 1A.

(B) Representative images of EdU-containing *myl7*:EGFP$^{twu34+}$ cardiomyocytes (arrows) at different time points after cryoinjury. A single intraperitoneal injection of EdU was performed 24 hours prior to harvesting hearts. Quantification of these data is found in Figure 1B.

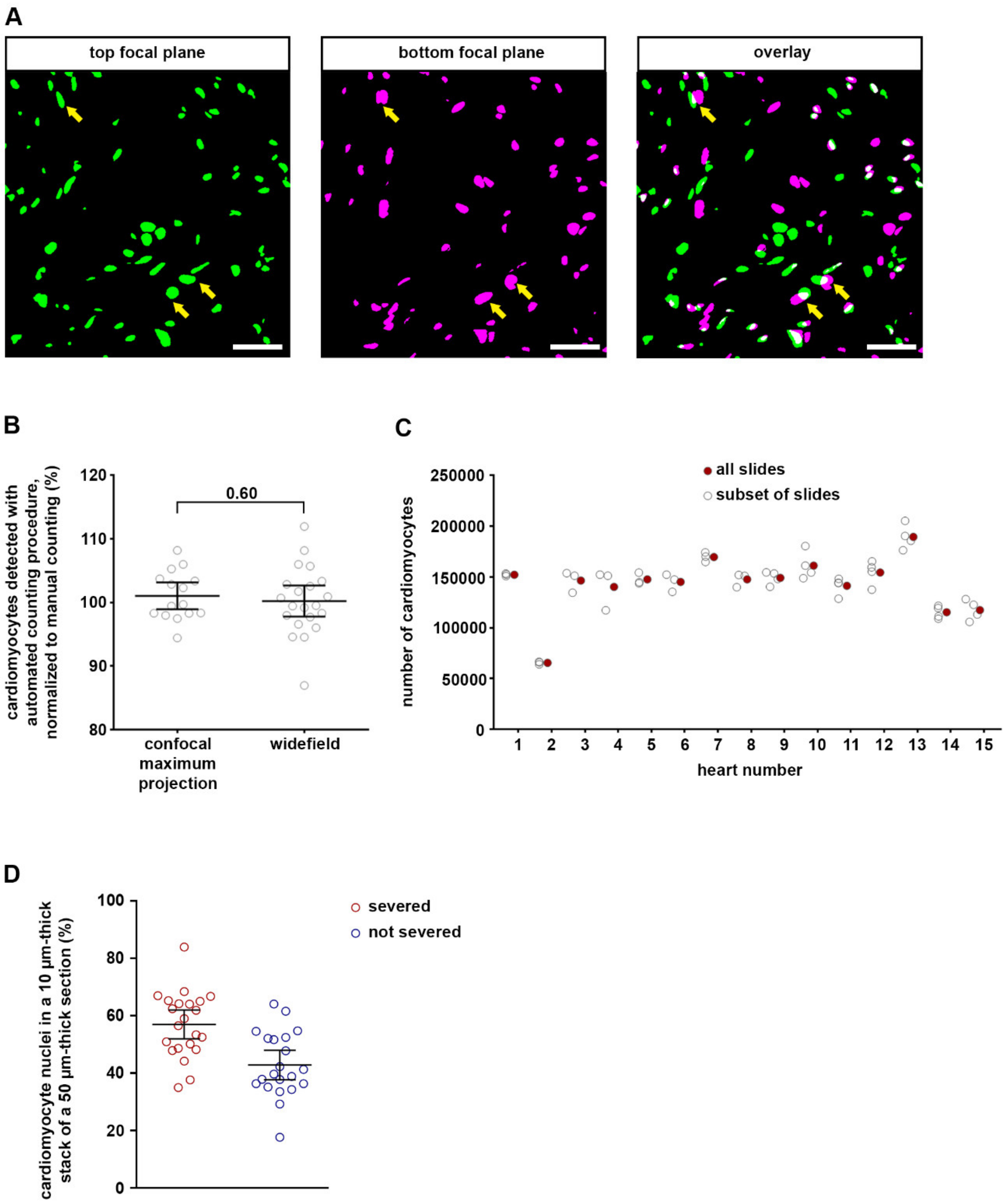


**Supplementary Figure 2. Validation of a method for counting cardiomyocyte numbers on cryosections.**

(A) The number of cardiomyocyte nuclei that are superimposed on maximum projection confocal stacks of 10 µm heart sections is low. Nuclei were outlined in the top (green) and bottom (magenta) optical plane of a confocal z-stack of 10 µm cryosections, 6-7 µm apart from each other, and overlap regions (white) were analyzed. Distinct overlapping cardiomyocyte nuclei are marked with yellow arrows. Scale bar, 100 µm.

(B) Automated counting using widefield microscopy images detects comparable numbers of cardiomyocytes as automatic or manual counting using confocal z-stack images. Error bars, CI 95%. n = 9 hearts, 1-3 sections per heart analyzed. Unpaired two-tail t-test.

(C) Subsampling does not significantly affect the estimation of the total number of cardiomyocytes. Comparison of the total number of cardiomyocytes per heart estimated from different subsets of slides, representing 1/6th of the sections of the whole ventricle, with the total number determined by counting all sections. n = 15 hearts.

(D) The number of cardiomyocyte nuclei was determined within a 10 µm thick confocal substack centered in the middle of a 50 µm thick cryosection. Nuclei that could be detected in the top or bottom z-plane of the substack and at the same time in the adjacent z-plane outside the sub-stack were considered "severed", that is they were cut by cryosectioning. On average, 57% of cardiomyocyte nuclei were severed in 10 µm-thick cryosections, and conversely 43% were not cut through, that is they could only be detected within the substack. Error bars, CI 95%. n = 6 hearts, 2-4 sections per heart analyzed.

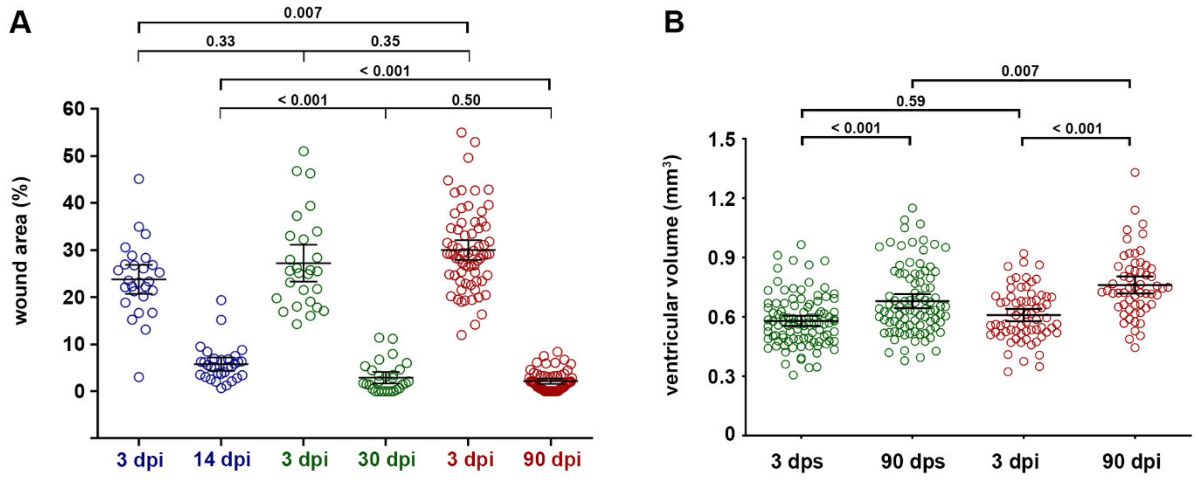


**Supplementary Figure 3. Data on wound area and ventricular volume.**

(A) Wound area at 3 dpi accounts for 25-30% of the ventricle area. The average wound area has dropped to 6%, 3% and 2% at 14, 30 and 90 dpi, respectively. Different colors indicate data from the three cohorts of fish that we injured, each with its own 3 dpi control. Error bars, CI 95%. n (hearts) = 27 (3 dpi 14-days group), 31 (14 dpi), 27 (3 dpi 30-days group), 29 (30 dpi), 68 (3 dpi 90-days group), 58 (90 dpi). One-way ANOVA + Dunnett's multiple comparisons test.

(B) Volume of the ventricular myocardium (ventricular volume including the wound area) recovers in injured hearts beyond levels observed in sham injured hearts within 90 days. Error bars, CI 95%. n (hearts) = 99 (3 dps), 92 (90 dps), 68 (3 dpi), 58 (90 dpi). One-way ANOVA + Tukey’s multiple comparisons test.

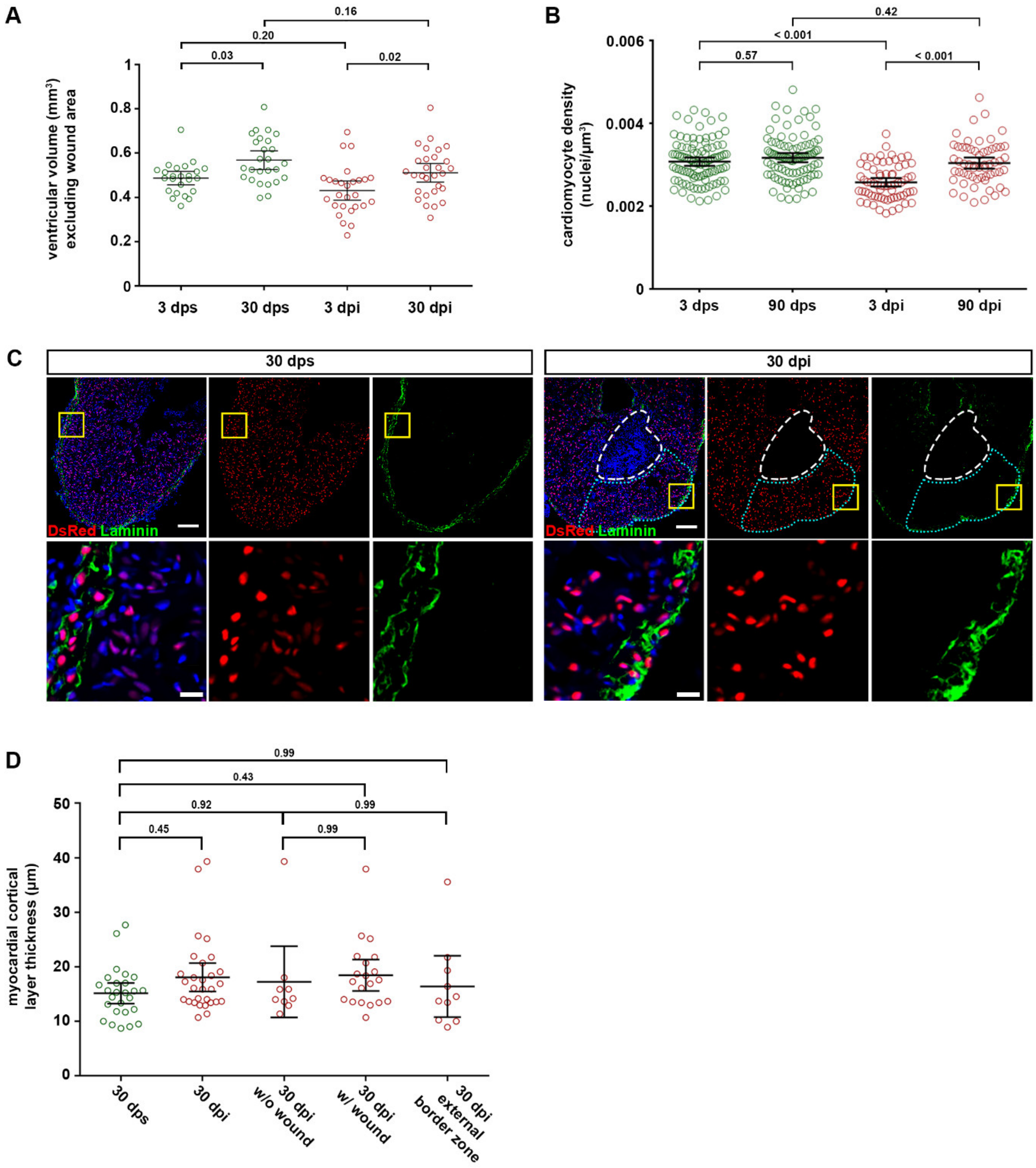


**Supplementary Figure 4. Overall myocardial morphology fully regenerates**

(A) Volume of the healthy myocardium (ventricular volume minus the wound area) recovers in injured hearts to levels similar to those of sham injured hearts within 30 days. Error bars, CI 95%. n (hearts) = 24 (3 dps), 26 (30 dps), 27 (3 dpi), 29 (30 dpi). One-way ANOVA + Tukey's multiple comparisons test. Observed difference 30 dpi vs 30 dps = 0.05 (10% of area in 30 dps); smallest significant difference = 0.07 (14% of 30 dps).

(B) Average cardiomyocyte density in the whole ventricle does not differ between injured fish at 90 dpi and sham-injured fish at 90 dps. Error bars, CI 95%. n (hearts) = 99 (3 dps), 92 (90 dps), 68 (3 dpi), 58 (90 dpi). One-way ANOVA + Tukey's multiple comparisons test. Observed difference 90 dpi vs 90 dps = 0.00013 (4% of density in 90 dps); smallest significant difference = 0.00026 (8% of 90 dps).

(C) Representative images of the compact cortical layer of cardiomyocytes, labeled by laminin immunostaining, at 30 days post sham-injury or cryoinjury. In the 30 dpi panel, dashed white lines indicate the remaining wound area and the dotted cyan outline marks the external wound border zone. Yellow rectangles mark the region in the top figures magnified in the bottom insets. Scale bars, 100 μm.

(D) Quantification of data shown in (C). Error bars, CI 95%. n (hearts) = 26 (30 dps), 29 (30 dpi), 9 (30 dpi no wound), 20 (30 dpi wound), 10 (30 dpi external border zone). One-way ANOVA + Tukey's multiple comparisons test.

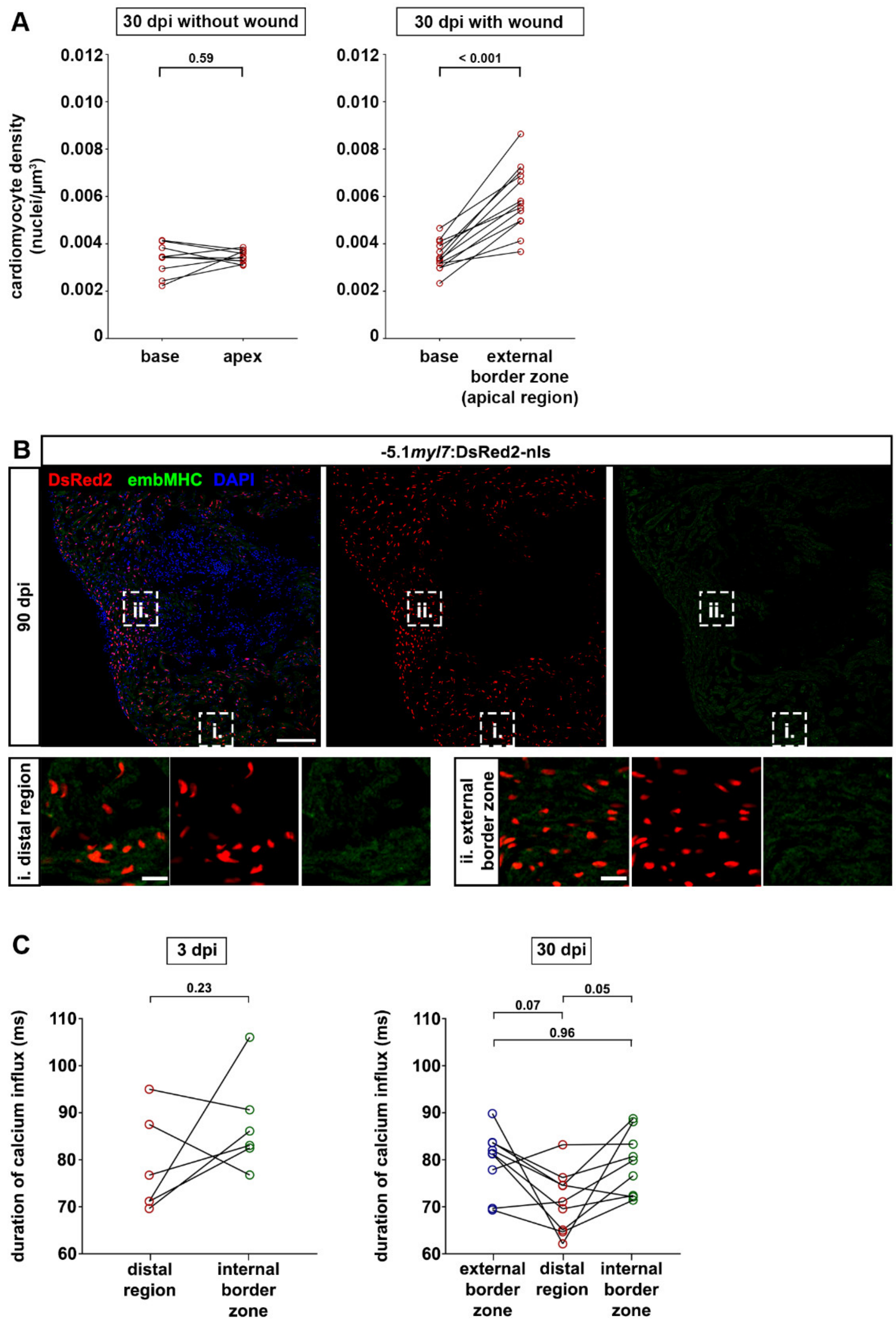


**Supplementary Figure 5. Data on cardiomyocyte density, embryonic marker expression and calcium influx in the external wound border zone**

(A) At 30 dpi, cardiomyocyte density at the base and the apex of the same ventricle does not differ in hearts without wound. In hearts with incomplete scar resorption, cardiomyocytes at the external wound border (apical region) have a higher density compared to cardiomyocytes at the base. n (hearts) = 9 (without wound), 13 (with wound). Paired two-tail t-test.

(B) In hearts with residual scars at 90 dpi, embMHC cannot be detected in cardiomyocytes either in distal regions of the myocardium (boxed area i.) nor at the external border zone (boxed area ii.). n = 0/16 hearts with embMHC$^{+}$ cardiomyocytes.

(C) Duration of calcium influx in the distal region and at the internal wound border zone of 3 dpi hearts (left panel) and in the distal region, internal and external wound border zone of 30 dpi hearts (right panel). Error bars, CI 95%. n (hearts) = 6 (3 dpi), 9 (30 dpi). Paired two-tail t-test.

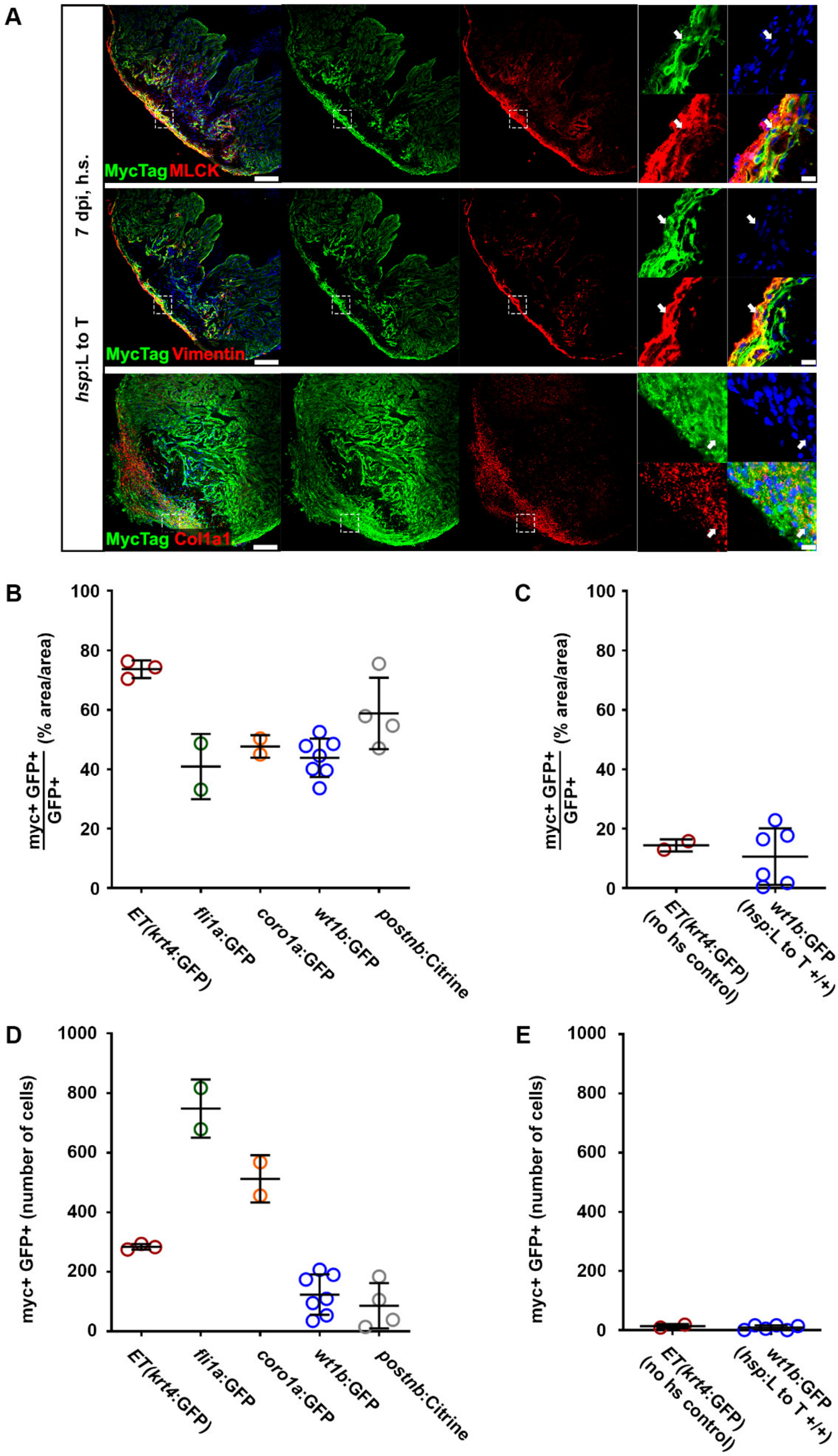


**Supplementary Figure 6. The *hsp*: L to T transgene is robustly expressed in all major cellular lineages of the adult heart**

(A) At 7 dpi, upon heat-shocks, luciferase-myc expression is detected in cells (arrows) positive for the fibroblast markers MLCK, Vimentin and Col1a1 in the ventricle of adult *hsp70l*:LOXP-luc-myc-

STOP-LOXP-dTomato, *cryaa*:YPet$^{ulm9}$ (*hsp*: L to T) non-recombined hearts. Boxed areas are shown in magnified view. Scale bars, 40 µm (overview), 5 µm (magnified view).

(B) Ratio of myc$^{+}$GFP$^{+}$ area over total GFP$^{+}$ area (in percentage) per section from the ventricle of adult *hsp*:L to T fish carrying in addition the indicated transgenic cell type reporters. Fish were not recombined, hearts injured, fish subjected to repeated heat-shocks and analyzed at 7 dpi. Data are shown for Et(*krt4*:eGFP$^{sqet27}$)$^{+}$ epicardial cells, *fli1a*:eGFP$^{y1Tg+}$ endocardial cells, *coro1a*:eGFP$^{hkz04tTg+}$ leukocytes, *wt1b*:EGFP$^{li1Tg+}$ epicardial cells, fibroblasts and macrophages, and *postnb*:Citrine$^{cn6Tg+}$ fibroblasts. Error bars, SD. n (hearts) = 3 (Et(*krt4*:eGFP)), 2 (*fli1a*:eGFP), 2 (*coro1a*:eGFP), 7 (*wt1b*:EGFP), 4 (*postnb*:Citrine).

(C) Background signals of myc staining shown as ratio of myc$^{+}$GFP$^{+}$ area over total GFP$^{+}$ area (in percentage) per section from the ventricle of adult *hsp*:L to T fish that were not recombined, and analyzed at 7 dpi. Double transgenics with Et(*krt4*:eGFP$^{sqet27}$)$^{+}$ (not heat-shocked) and *wt1b*:EGFP$^{li1Tg+}$ single transgenics were analyzed. Error bars, SD. n (hearts) = 2 (Et(*krt4*:eGFP), no hs control), 6 (*wt1b*:EGFP, no *hsp*: L to T).

(D) Absolute number of myc$^{+}$GFP$^{+}$ cells per ventricular section for the same double transgenic heart samples as shown in (B). Error bars, SD. n (hearts) = 3 (Et(*krt4*:eGFP)), 2 (*fli1a*:eGFP), 2 (*coro1a*:eGFP), 7 (*wt1b*:EGFP), 4 (*postnb*:Citrine).

(E) Absolute number of myc$^{+}$GFP$^{+}$ cells per ventricular section for the same control heart samples as shown in (C). Error bars, SD. n (hearts) = 2 (Et(*krt4*:eGFP), no hs control), 6 (*wt1b*:EGFP, no *hsp*: L to T).

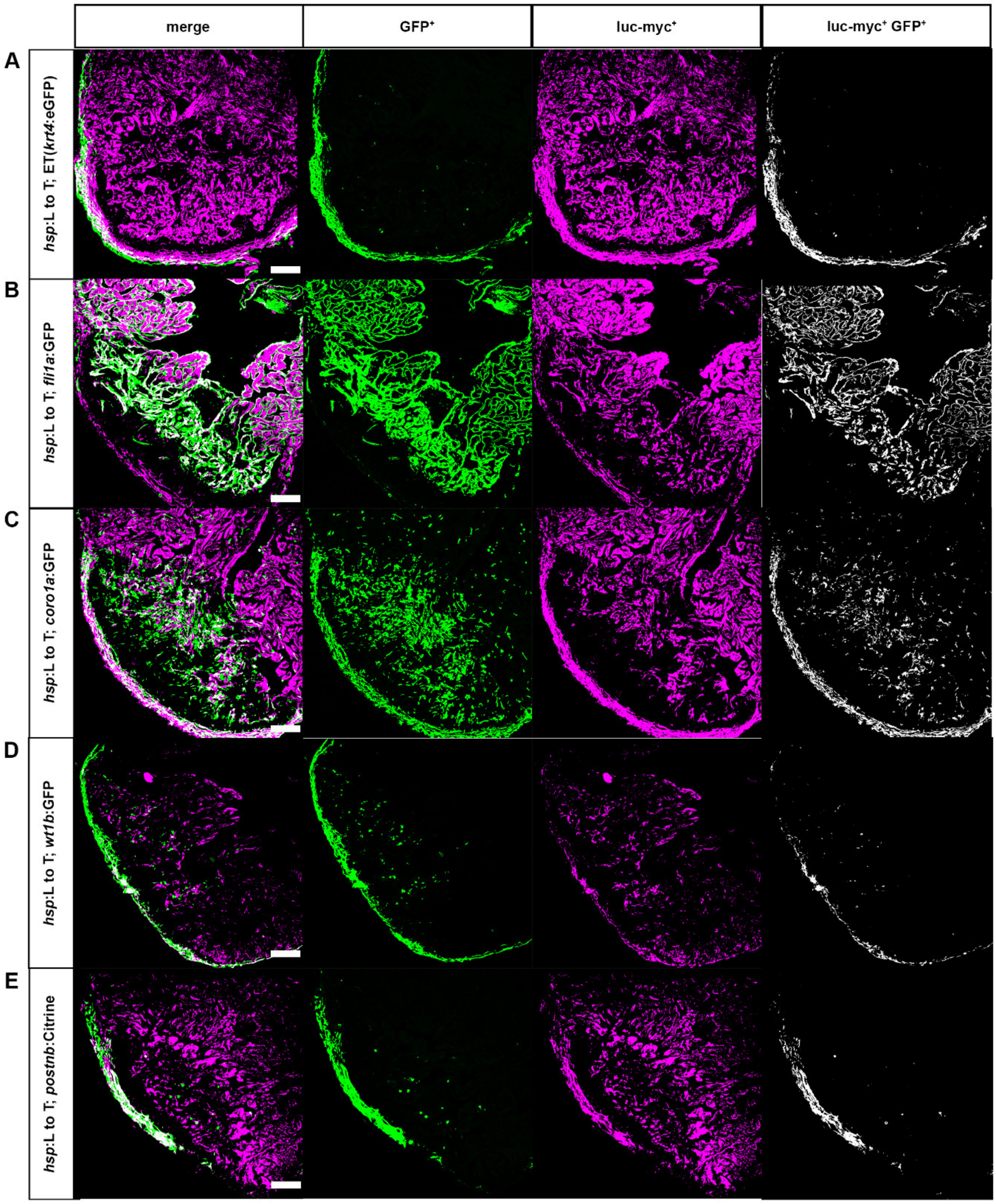


**Supplementary Figure 7. Spatial distribution of the *hsp*: L to T transgene in non-myocyte cellular lineages of the adult heart**

At 7 dpi, in heat-shocked fish, myc-tagged luciferase expression is broadly distributed in non-myocytes of the ventricle of adult *hsp70l*:LOXP-luc-myc-STOP-LOXP-dTomato, *cryaa*:YPet$^{ulm9}$ (*hsp*:L to T) hearts, in particular in the area occupied by Et(*krt4*:eGFP$^{sqet27}$)$^{+}$ epicardial cells (A), *fli1a*:eGFP$^{y1Tg+}$ endocardial cells (B), *coro1a*:eGFP$^{hkz04tTg+}$ leukocytes (C), in *wt1b*:EGFP$^{li1Tg+}$ epicardial cells, fibroblasts and macrophages (D), and in *postnb*:Citrine$^{cn6Tg+}$ fibroblasts (E). Saturated lookup tables are shown for GFP$^{+}$ area (green), luc-myc$^{+}$ area (magenta) and luc-myc$^{+}$ GFP$^{+}$ double positive area (white). Scale bars, 50 µm.

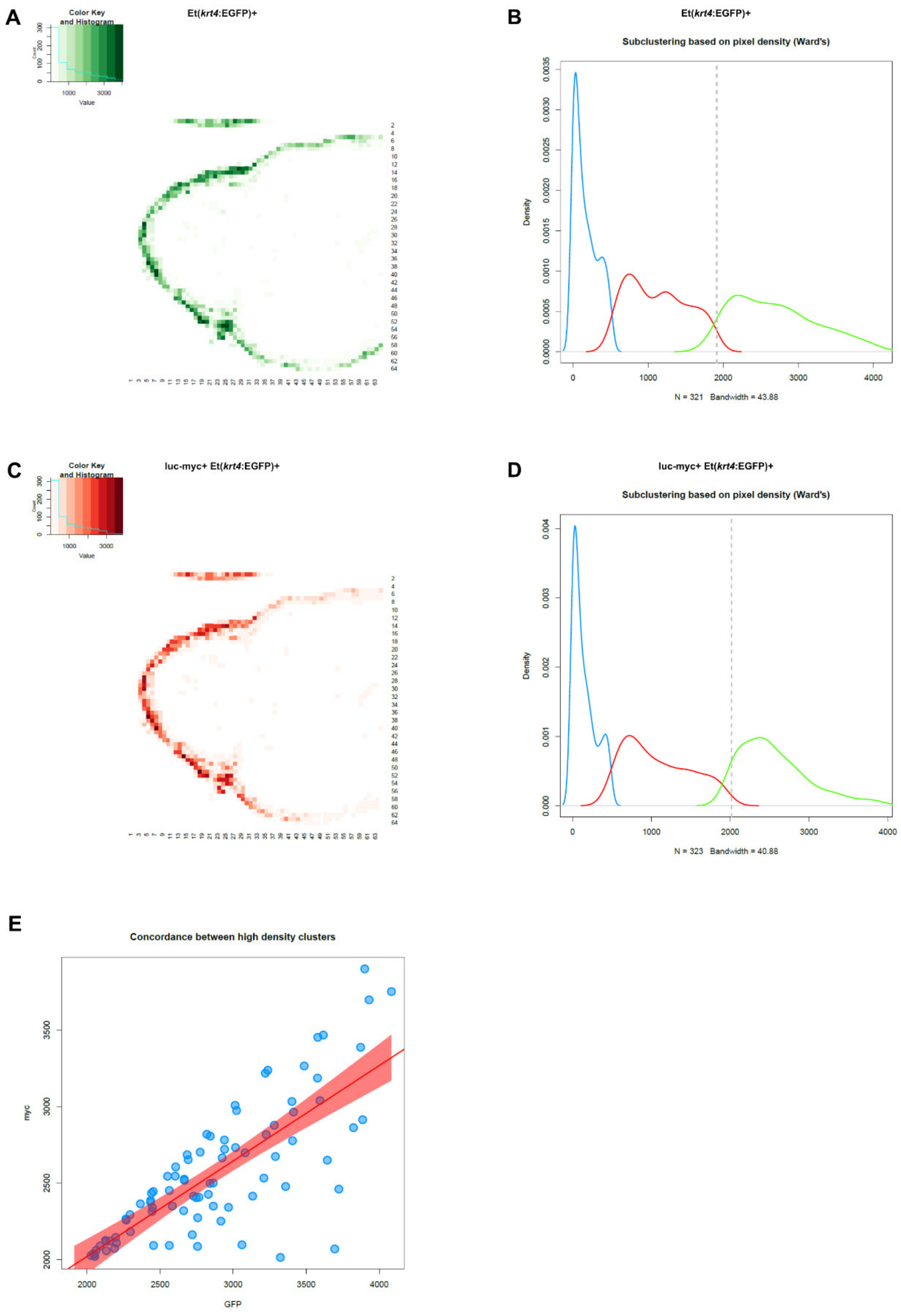


**Supplementary Figure 8. Signal concordance analysis of the *hsp*: L to T transgene and GFP expression in the epicardium of the adult heart**

Shown here are exemplary images of the workflow used to perform concordance analysis between luc-myc expression in the *hsp*: L to T tracer transgene and other transgenes labelling the epicardium, endocardium and macrophages and neutrophils.

(A) Binned quantitative plots of the GFP$^{+}$ area in a section from the ventricle of a heat-shocked adult *hsp*:L to T, Et(*krt4*:eGFP$^{sqet27}$) heart at 7 dpi. Upper left corner, histogram representing the pixel density of the binned clusters vs. the count of binned clusters for each pixel density.

(B) Subclustering based on pixel density of the binned image shown in (A) identifies three clusters of low (blue), intermediate (red), and high (green) pixel count regions. X-axis represents ranges of intensities (no. of pixels per bin) while y-axis is their density distribution. The dashed grey line represents the threshold used for the cut-off of high density clusters.

(C) Binned quantitative plots of the luc-myc$^+$ GFP$^+$ area of the same section shown in (A). Upper left corner, histogram representing the pixel density of the binned clusters vs. the count of binned clusters for each pixel density.

(D) Subclustering based on pixel density of the binned image shown in (C) identifies three clusters of low (blue), intermediate (red), and high (green) pixel count regions. X-axis represents ranges of intensities (no. of pixels per bin) while y-axis is their density distribution. The dashed grey line represents the threshold used for the cut-off of high density clusters.

(E) A plot depicting statistically significant concordance between GFP$^+$ (x-axis) and luc-myc$^+$ GFP$^+$ (y-axis) high density clusters identified with the green curves of (B) and (D), respectively. Shown is also a linear regression trendline and its corresponding 95% confidence interval.

A

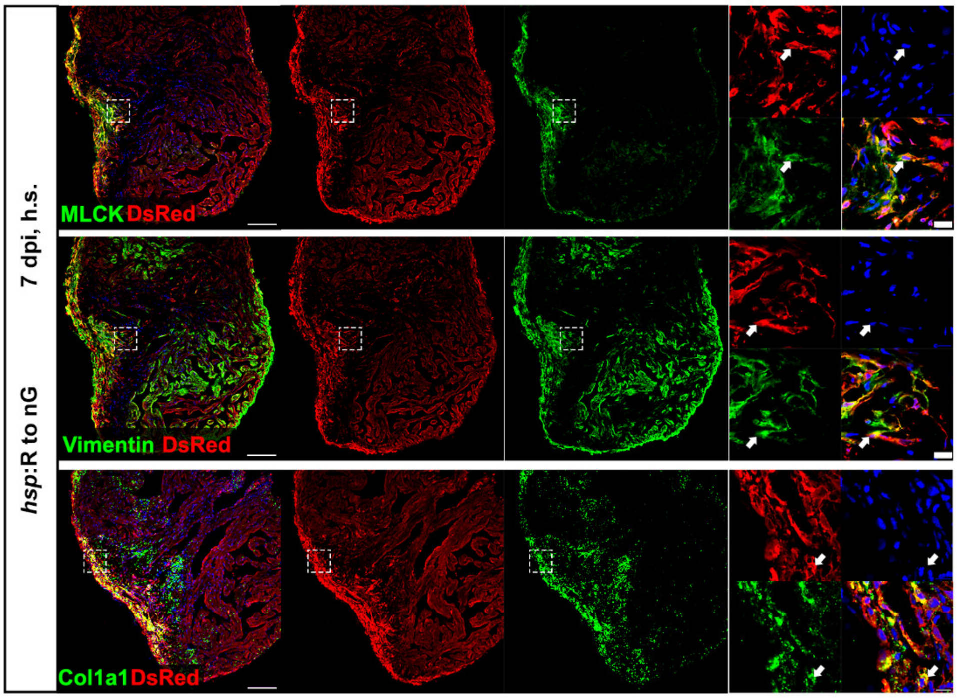


B

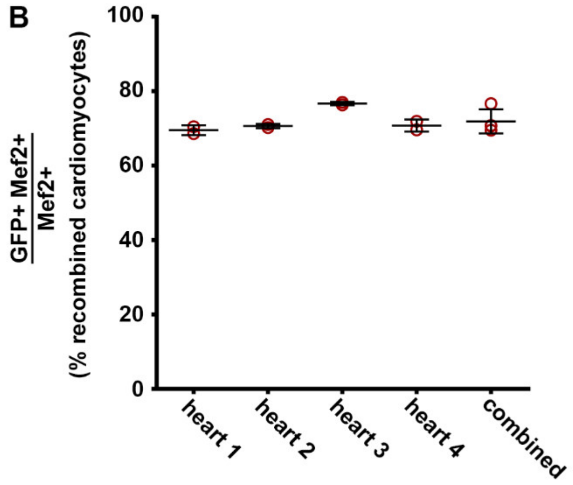


**Supplementary Figure 9. The *hsp*:R to nG transgene is expressed in fibroblasts in the juvenile heart and high recombination efficiency can be achieved by embryonic Tamoxifen treatment.**

(A) At 7 dpi, upon heat-shocks, DsRed expression is detected in cells (arrows) positive for the fibroblast markers MLCK, Vimentin and Col1a1 in the ventricle of non-recombined juvenile *hsp*:R to nG$^{tud9}$ hearts. Boxed areas are shown in magnified view. Scale bars, 40 µm (overview), 5 µm (magnified view).

(B) Recombination efficiency expressed as number of EGFP$^{+}$Mef2$^{+}$ cardiomyocytes relative to the total number of Mef2$^{+}$ cardiomyocytes in juvenile *cryaa*:DsRed,-5.1*myl7*:Cre-ERT2$^{pd10}$; *hsp70l:*LOXP-DsRed2-LOXP-NLS-EGFP$^{tud9}$ (*hsp*:R to nG) hearts upon heat shock that were subjected to 4-OH Tamoxifen treatment at embryonic stages as indicated in Figure 7E.

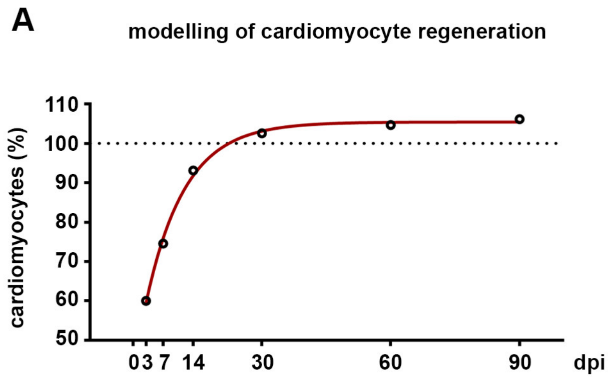


**Supplementary Figure 10: A theoretical model of cardiomyocyte proliferation dynamics during regeneration.**

(A) Modelling of the time-course of cardiomyocyte regeneration in response to injury that kills 38% of cardiomyocytes using the following assumptions: (1) all cardiomyocytes located at the wound border that are PCNA$^+$ at a given time-point also proliferate; (2) cell cycle length is 24 hours and remains constant throughout regeneration. Modelling was based on the PCNA data shown in Figure 1A. That is, the rate of PCNA$^+$ cardiomyocytes at 3, 7, 14, 30 and 60 dpi was used as given in the figure, while the rate for the days for which no measurements were done were extrapolated as described in the Extended Materials & Methods section, e.g. the rate for days 8 to 13 was the average of the available 7 dpi and 14 dpi data.

### Supplementary Table

**Supplementary Table 1.** Summary statistics of the non-parametric Kendall's concordance analysis between GFP and myc signals. The Kendall's coefficient is given in the column 'estimate' (ranging between 0 and 1). The following additional columns are also provided: associated degrees of freedom (df), the chi-square statistic (statistic), test of significance based on p-value (p.value), the level of significance (Sig.; p.value < 0.001 = ***, p.value < 0.01 = **, p.value < 0.05 = *, and NS or Not Significant), and the conclusion based on the p-value (Concordance = significant p-value; No concordance = NS). See methods for further details on the statistical methodology.

| Image File | estimate | df | statistic | p.value | Sig. | Concordance |
|---|---|---|---|---|---|---|
| coro1a.heart1 | 0.748 | 263 | 393.443 | 3.20E-07 | *** | Concordance |
| coro1a.heart2 | 0.824 | 145 | 238.876 | 1.48E-06 | *** | Concordance |
| et27.heart1 | 0.666 | 76 | 101.250 | 0.028 | * | Concordance |
| et27.heart2 | 0.612 | 56 | 68.566 | 0.121 | NS | No Concordance |
| et27.heart3 | 0.874 | 84 | 146.754 | 2.73E-05 | *** | Concordance |
| fli1a.heart1 | 0.628 | 187 | 234.893 | 0.010 | * | Concordance |
| fli1a.heart2 | 0.527 | 252 | 265.385 | 0.269 | NS | No Concordance |
| wt1b.heart1 | 0.652 | 64 | 83.408 | 0.052 | NS | No Concordance |
| wt1b.heart2 | 0.738 | 27 | 39.829 | 0.053 | NS | No Concordance |
| wt1b.heart3 | 0.491 | 24 | 23.584 | 0.486 | NS | No Concordance |
| wt1b.heart4 | 0.620 | 49 | 60.772 | 0.121 | NS | No Concordance |
| wt1b.heart5 | 0.811 | 12 | 19.461 | 0.078 | NS | No Concordance |
| wt1b.heart6 | 0.400 | 14 | 11.200 | 0.670 | NS | No Concordance |
| wt1b.heart7 | 0.566 | 46 | 52.052 | 0.250 | NS | No Concordance |
